# Sick of Eating: eco-evo-immuno dynamics of predators and their trophically acquired parasites

**DOI:** 10.1101/2020.05.26.117622

**Authors:** Samuel R. Fleischer, Daniel I. Bolnick, Sebastian J. Schreiber

## Abstract

When predators consume prey, they risk becoming infected with their prey’s parasites, which can then establish the predator as a secondary host. For example, stickleback in northern temperate lakes consume benthic or limnetic prey, which are intermediate hosts for distinct species of parasites (e.g. *Eustrongylides* nematodes in benthic oligocheates and *Schistocephalus solidus* copepods in limnetic copepods). These worms then establish the stickleback as a secondary host and can cause behavioral changes linked to increased predation by birds. In this study, we use a quantitative genetics framework to consider the simultaneous eco-evolutionary dynamics of predator ecomorphology and predator immunity when alternative prey may confer different parasite exposures. When evolutionary tradeoffs are sufficiently weak, predator ecomorphology and immunity are correclated among populations, potentially generating a negative correlation between parasite intake and infection.

## Introduction

Many predators acquire parasites by consuming infected prey (Iritani and Sato 2018; Rogawa et al. 2018). For instance, in a food web in the Carpinteria Salt Marsh, there were twice as many predator-parasite links than predator-prey links (Lafferty et al. 2006). Because of the ubiquity of trophically-transmitted parasites, community ecologists are increasingly interested in the impact of parasites on communities (Anderson and Sukhdeo 2011; Sukhdeo 2012; Wood and Johnson 2015). For example, parasites have been shown to alter both food web dynamics and structure (Cortez and Weitz 2014; Lafferty et al. 2006; Velzen and Gaedke 2017). Recent food web models are thus incorporating the effects of diet-driven infection rates, which depend on a host’s position within the food web. However, after a parasite passes through this ‘encounter filter,’ it must then pass through a ‘compatibility filter’ by establishing an infection despite the host’s immune response. Therefore, to understand the role of parasitism within an ecological community we must consider both the host diet (encounters) and host immunity (compatibility).

Recent empirical and theoretical studies on eco-evolutionary feedbacks can help us understand how predators can encounter different trophically-transmitted parasites. In particular, predator-prey interactions may drive trait evolution in the prey to facilitate predator evasion or in the predator to optimize foraging success. Yoshida et al. (2003) found that prey evolution can affect the period and phase difference in predator-prey cycles, and Becks et al. (2010) found that the amount of heritable variation in a prey defense trait can shift dynamics between equilibrium and oscillatory states. These example illustrate how trait evolution modifies predator-prey interactions in ways that can change their population dynamics, coexistence, food web structure, and, hence, the rate of predator enounter with trophically-transmitted parasites.

For trophically transmitted parasites, eco-evolutionary dynamics can lead to changing risk of parasite infection through time. Likewise exposure risk can differ between ecologically divergent consumer individuals or populations. A well-studied species that exhibits this phenomenon is the threespine stickleback (*Gasterosteus aculeatus*), a small fish found in north temperate coastal habitats. Stickleback populations in different lakes typically specialize on eating the local abundant prey and evolve functional morphology to suit this niche (Lavin and McPhail 1986, 1985). In lakes with multiple abundant prey, stickleback are usually generalists, but individuals’ ecomorphology is correlated with their diet (Snowberg et al. 2015). The foraging diversity within and among stickleback populations affects parasite exposure. These associations between ecomorphology, diet, and infection risk have also been found in coastal California sea otters (Estes et al. 2003; Johnson et al. 2009; Tinker et al. 2008), cichlids (Hayward et al. 2017), and barbs (Sibbing et al. 1998).

Because of the complex lifecycle of the trophically-transmitted cestodes and nematodes that infect stickleback, only individuals who consume particular prey are at risk of infection by those parasites. The cestode *Schistocephalus solidus* uses cyclopoid copepods as a first host, which are eaten by stickleback in open water (limnetic habitat). In contrast, the ne-matode *Eustronglyides sp.* uses benthic-dwelling oligocheate worms as their primary host. As a result, stickleback diet should be correlated with parasite intake rates: individuals consuming more limnetic copepods should have higher *S. solidus* infection rates. This expectation has been confirmed within each of many populations showing correlations between individuals’ ecomorphology and infection status (Stutz et al. 2014). One might expect this correlation between stickleback ecomorphology and infection risk to hold not only within populations, but also among populations. However, Stutz et al. (2014) found that nematode prevalence was lowest in populations where we would a priori expect it to be highest (where fish consume more benthic prey). Stutz et al. (2014) suggested that correlated evolution of diet and immunity might be the cause of this negative correlation. And, since these ces-todes and nematodes come from different prey and are phylogenetically distant parasites, it seems possible that immunity to infection might need to be specialized to one or the other. Motivated by this unexpected negative correlation, we seek to address three questions.

*First, under what conditions is there a reciprocal feedback between niche and immune evolution? If these conditions do not hold, when does the evolution of the predator’s trophic niche determine the evolution of its immune system, or vice versa?* Environmental conditions may dictate the relative availability of alternative prey, which in turn generates selection on the predator’s foraging morphology. The subsequent eco-evolutionary feedbacks involve coupled changes in species abundances and predator traits. Changes in both prey availability and predator efficiency will alter the predator’s diet and thus change its exposure to parasites, which will likely drive evolution of the predator’s immune system to resist whichever parasites represent the greatest risk. On the other hand, the evolution of a predator’s immune trait can reduce the harmful effects of parasites, enabling niche expansion or a niche shift onto a formerly hazardous prey. Trophic traits may then subsequently evolve to optimize attack efficiency on this new diet. When do these effects create a feedback loop and cause multivariate-trait evolutionary cycles?

*Second, how are trophic and immune trait values correlated?* Even if traits are genetically independent, their selection pressures may not be. The predator’s immune trait may affect the selection pressure on morphology, and vice versa. The joint evolution of traits may lead to correlations between diet and immunity either because habitat differences favor bivariate outcomes or because populations are at different phases of some cyclical dynamic.

*Third, when does the joint evolution of niche and immune traits, along with predator-prey dynamics, obscure the relationship between parasite exposure risk and actual infection rates?* As hypothesized by Stutz et al (2014), evolution of predator immune traits may negate or even reverse an expected positive correlation between intake of, and infection by, a particular parasite. Thus, populations frequently exposed to particular parasites may have lower infection rates than populations that are rarely exposed (and hence susceptible) to the few parasites they do encounter.

## Models

Let *P* = *P* (*t*) be the density of a predator population and *N*_*i*_ = *N*_*i*_(*t*) be the densities of prey populations *i* for *i* = 1, 2. Each prey species experiences logistic growth in the absence of the predator, with intrinsic growth rates *r*_*i*_ and carrying capacities *K*_*i*_. The predator species has a per-capita death rate *d*, attacks prey species *i* with attack rate *a*_*i*_, and converts food into offspring with efficiency *b*_*i*_.

A percentage *c*_*i*_ of prey *i* individuals are infected with parasite *i* that decreases predator fecundity by *m*_*i*_*S*_*i*_, where *m*_*i*_ is the maximal negative effect of parasite *i* and *S*_*i*_ ≤ 1 is a measure of predator susceptibility to parasite *i*. Low *S*_*i*_ corresponds to a strong immunity to parasite *i*. Then the ecological dynamics are given by

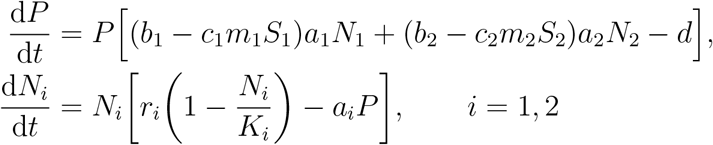

The attack rate of the predator on prey *i* is determined by a quantitative trait *x*. Attack rate of prey species *i* is maximal when *x* = *θ*_*i*_, where *θ*_*i*_ is the optimum trait to consume prey *i*, and decreases in a Gaussian manner as |*x* − *θ*_*i*_| increases (as in Schreiber et al. (2011)). Specifically, the attack rate *a*_*i*_(*x*) on prey *i* equals

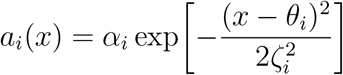

where *α*_*i*_ is the maximal successful attack rate on prey *i*, and *ζ*_*i*_ is the width of the attack rate function. The smaller *ζ*_*i*_, the more phenotypically specialized a predator must be to use prey *i*. Thus, predator populations with greater *ζ*_*i*_ values experience less pressure to evolve morphological specialization.

The susceptibility of the predator to infection by parasite *i* is determined by a quantitative trait *y*. Susceptibility is minimized when *y* = *ϕ*_*i*_, where *ϕ*_*i*_ is the optimum trait to resist parasite *i*, and increases in a Gaussian manner as |*y* − *ϕ*_*i*_| increases. Specicically, the susceptibility *S*_*i*_(*y*) to parasite *i* equals

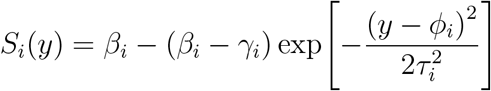

where *β*_*i*_ ≤ 1 and *γ*_*i*_ < *β*_*i*_ are the maximum and minimum susceptibility to parasite *i*, respectively, and *τ*_*i*_ is the width of the immunity function. The smaller *τ*_*i*_, the more immunologically specialized a predator must be to significantly reduce susceptibility to infection by parasite *i*. Thus, predator populations with greater *τ*_*i*_ values experience less pressure to evolve immunological specialization. We have in mind a model of constitutively expressed innate immunity rather than adaptive immunity that is induced and grows following initial exposure.

The per-capita growth rate *W* (fitness) of a predator with ecomorphology *x* and immunity *y* is given by

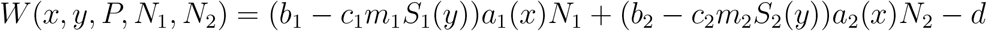

and the per-capita growth rate *Y*_*i*_ of prey interacting with predators with ecomorphology *x* is given by

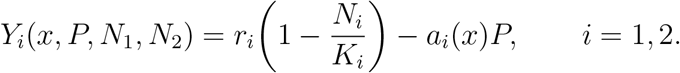

We assume the predator traits *x* and *y* are genetically independent and normally distributed over the population with means 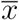 and 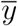, respectively. Let 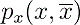 and 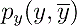 denote these distributions:

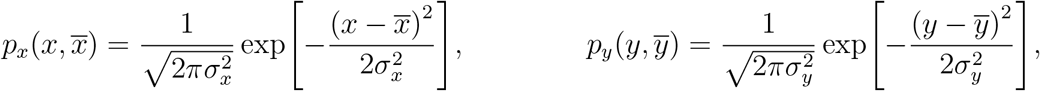

where 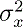 and 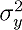 are the total phenotypic variances of traits *x* and *y*, respectively. Let 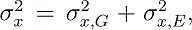 where 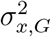 is the phenotypic variation of trait *x* due to genotype and 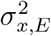 is the phenotypic variation of trait *x* due to environmental conditions. Similarly, let 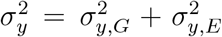. Here, we omit genetic-by-environmental interactions for mathematical simplicity, though these are common for both trophic and immunological traits. Note that the environmental variance component is not adaptive placticity (e.g. not directionally dictated by prey availability or parasite exposure experience).

Integrating across the predator distribution of phenotypes, the average per-capita growth rate of the predator population 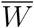 equals

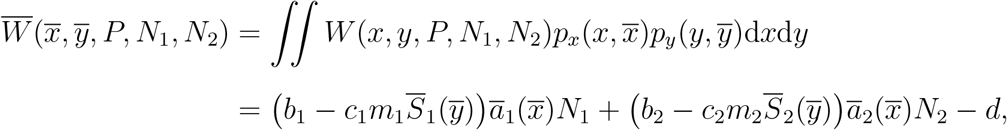

where 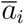 and 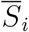 are the averaged attack rate and susceptibility:

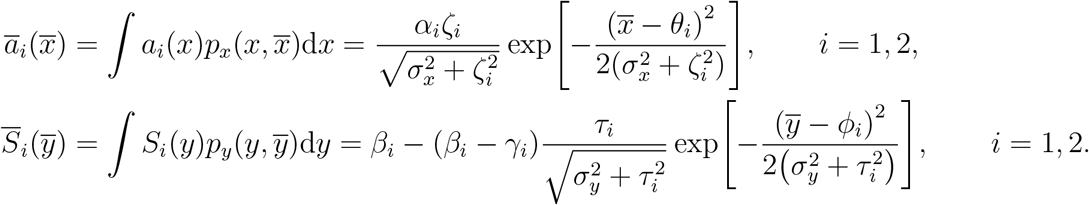

The average per-capita growth rate 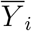 of the prey *i* population is given by

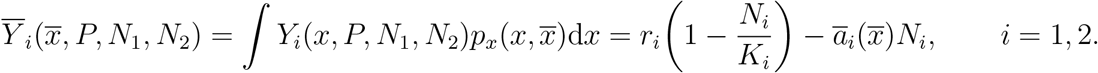

These functions describe the ecological dynamics:

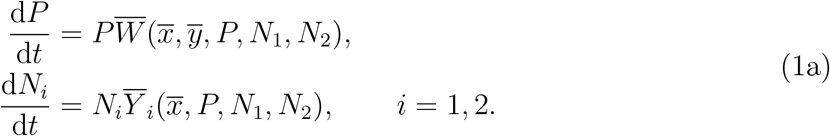

See Appendix A for additional details regarding the model formulation.

Provided the predator trait distributions *p*_*x*_ and *p*_*y*_ stay normal with constant variance over time, Lande (1976) showed that the rates of change of the average traits are proportional to the derivative of the average fitness 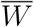 with respect to that trait. The constants of proportionality are the portions of phenotypic variance due to genetic variation. We assume the morphological and immunological traits are genetically independent, and thus the evolutionary dynamics of 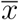 and 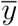 are given by:

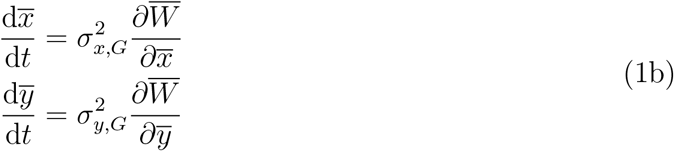

where

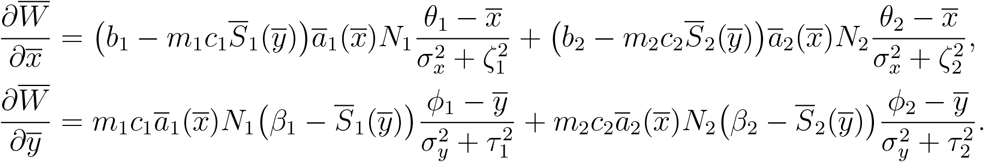

## Methods

We use four numerical and analytical approaches to explore the three questions posed in the introduction: (i) numerical simulations of population and evolutionary dynamics, (ii) analytical results obtained in the limit of slow evolution (low heritability) and timescale differences between the evolution of the two traits, (iii) Latin hypercube sampling across parameter space to analyze the effect of model parameters on simulation outcomes, and (iv) numerical approximations of Lyapnuov exponents to determine conditions for stable or chaotic dynamical behavior.

### Numerical Simulations

We used standard numerical integration techniques (Runge-Kutta 4(5) with adaptive step size using Python’s `scipy.integrate.odeint` (Jones et al. 2001–)) to simulate the models (1a,1b) and (2). Parameters are given in Appendix B.

Lyapunov exponents describe how nearby trajectories behave in relation to a reference trajectory (Sprott 2003). Given an initial condition, a positive (negative) Lypunov exponent means that nearby trajectories on average move away from (towards) the reference trajectory, indicating chaos (stability) (Appendix C). We calculated Lyapunov exponents for full-model simulations over a two-dimensional subset of parameter space (*σ*_*y,G*_ vs. *τ*) to determine how the evolution of the immune trait 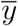 affects the eco-evolutionary dynamics.

### Analytic Reductions for Slow Evolution Dynamics

When the trait dynamics evolve at a sufficiently slower time scale than ecological dynamics, we can reduce the five-dimensional system to a two-dimensional system. This occurs, for example, if ecomorphological and immunity traits are only marginally heritable (small 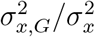 and 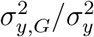). Then 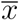 and 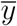 are effectively constant with respect to the changing population densities *P* and *N*_*i*_, *i* = *L, B*. In which case, the population dynamics of the fast ecological system converges to a unique globally stable attractor with a time-averaged ecological state 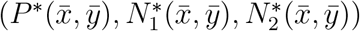 equal to the coexistence equilibrium 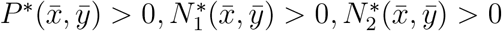 (Appendix D). Thus, on the evolutionary timescale, the trait dynamics are

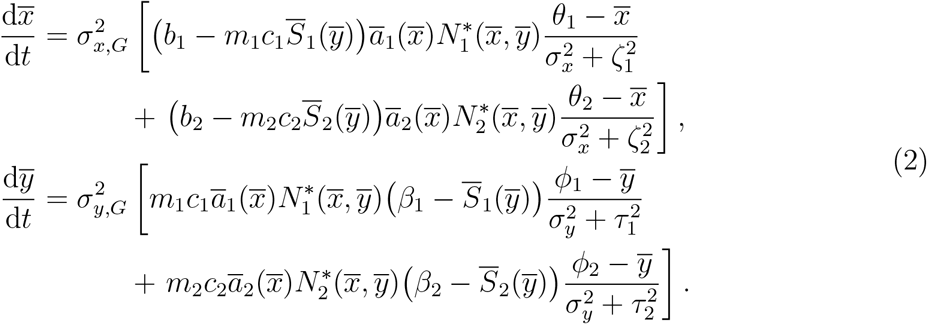

For the reduced system (2), we calculate nullclines, stable and unstable equilibria, and separatrices across a range of foraging trade-offs.

Beyond the separation of timescale between ecological and evolutionary dynamics, the niche and immune traits can themselves evolve on different timescales. As the two traits are genetically independent, one trait may evolve on a slower timescale than the other if the two traits differ significantly in their genotypic variance or their selection pressure. These differences can arise in three ways, as discussed in the Results section.

### Latin Hypercube Sampling

Latin hypercube sampling is a systematic way to explore high-dimensional parameter spaces to gain insight into how particular parameters correlate with certain outcomes, how parameters interact, and how outcomes correlate with each other (Stein 1987). In this study, we are interested in how ecological parameters (e.g., prey carrying capacities, parasite pathogenicity, etc.) determine the evolution of the predator’s foraging and immune traits. Using the equilibria of the slow-evolution model, we calculated the relative intake rates of the two prey types 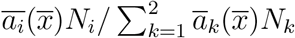, (*i* = 1, 2), the relative exposure rates to the two parasite types, 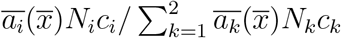, (*i* = 1, 2), as well as the relative parasite infection rates (Figure 1b) of the two parasite types 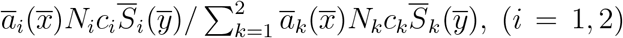, over a two-dimensional range of foraging tradeoffs (*ζ*_*i*_) and immune tradeoffs (*τ*_*i*_). For each tradeoff pair, we ran 4,000 simulations using Latin hypercube sampling, varying lake size (proxied by the ratio of limnetic to benthic carrying capacities *K*_1_*/K*_2_), limnetic and benthic maximal attack rates (*α*_1_, *α*_2_), limnetic and benthic parasite frequency in prey (*c*_1_, *c*_2_), limnetic and benthic parasitic effects on stickleback (*m*_1_, *m*_2_), limnetic and benthic prey growth rates (*r*_1_, *r*_2_), and initial stickleback ecomorphology and immunity 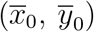. For each set of parameter values, we ran the two-timescale model (2) until 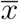 and 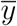 reached an evolutionary equilibrium and calculated the relative intake, exposure, and infection rates. We then plotted the results to gain insight about the joint evolution of niche and immune traits and how their evolution affects the relationship between diet, parasite exposure, and infection.

**Figure 1:**
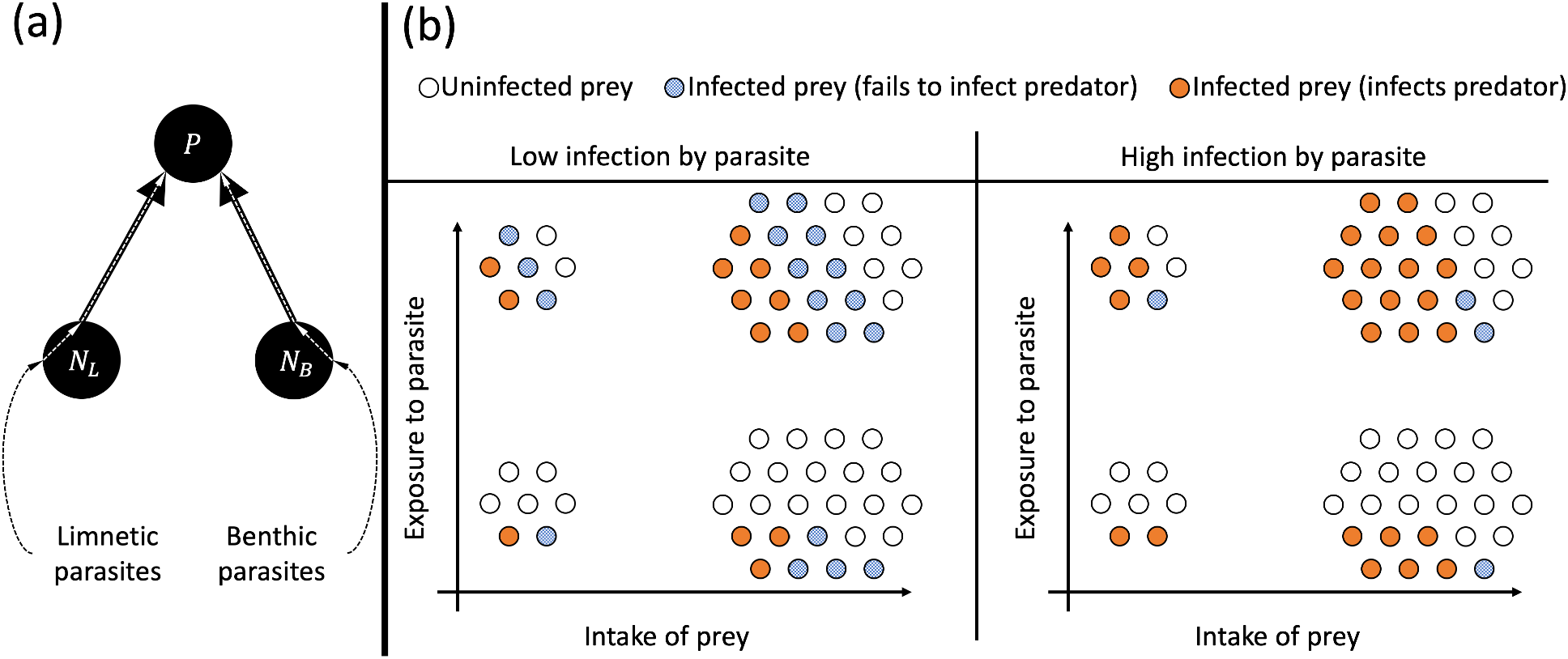
(a) Schematic of the model. The predator is exposed to limnetic and benthic parasites via intake of limnetic and benthic prey. The proportion of prey infected by parasites stays constant. The predator average morphology 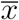 and average immune response 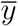 evolves in response to selection pressures caused by prey availability and parasite infection. Intake of prey 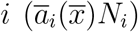 describes total intake of both infected and uninfected prey. Exposure to parasite 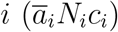 describes total intake of parasites (a constant proportion *c*_*i*_ of prey *i* are infected with parasites). Infection by parasite 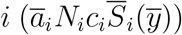 describes the total number of ingested parasites which successfully infect the predator (a proportion 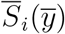 of ingested parasites infect the predator).

## Results

We first present results of the slow-evolution models to address how the evolution of niche affects the evolution of immunity and vice versa. We then present the Latin hypercube sampling results to address how the two traits are correlated across populations, as well as how that correlation affects the relationship between diet and infection across populations.

We conclude with a “within populations” perspective by using the full single-timescale five-dimensional model to examine temporal correlations in traits and diet and infection rates for systems with cyclic or chaotic dynamics.

### Three-timescale dynamics

For a given immune state 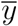, the average predator fitness 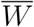 is unimodal with respect to the foraging trait 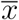 if

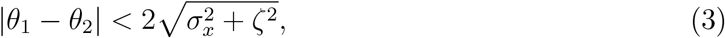

where *ζ* := *ζ*_1_ = *ζ*_2_ (Schreiber et al. 2011; Schreiber and Patel 2015). Namely, if foraging tradeoffs are weak relative to the phenotypic variation in foraging, then there is a single fitness maximum with respect to *x̄*. When the contributions of the two prey populations to predator fitness are equal (i.e. 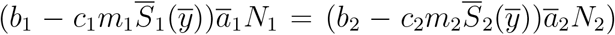), condition (3) is necessary and sufficient, but when prey contributions to predator fitness are unequal, predator fitness may be unimodal with respect to 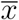 even if (3) does not hold.

Similarly, for a given foraging trait 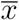, 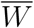 is unimodal with respect to the immune trait 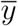 if the immune tradeoff is weak relative to the phenotypic variance in immunity, i.e.

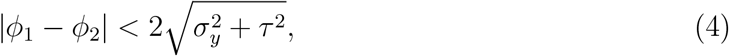

where *τ* := *τ*_1_ = *τ*_2_ (Appendix E). This condition is necessary and sufficient only when the difference between the effects of parasites infecting predators maximimally and minimally susceptible to those parasites is symmetric (i.e. 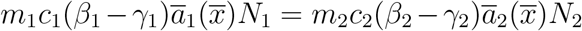). Again, if these differences are unequal, 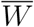 may still be unimodal with respect to 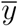 even if (4) does not hold.

There are three ways in which the predator traits may evolve at different timescales. First, all else being equal, the trait with a higher genotypic variance evolves more quickly than the other (Figure 2a,b,d,e,g,h,j,k). Second, if the parasite is rare or has a weak effect on the predator, then selection pressure on the immune trait 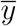 is weak and therefore evolves much slower than the foraging trait 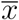 (Figure 2c,f,i,l). Third, weak tradeoffs in either trait result in weak selection pressure on that trait. In particular, large *ζ*_*i*_ corresponds to slower 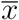 evolution (Figure 2d-f,j-l), and large *τ*_*i*_ corresponds to slower 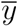 evolution (Figure 2a-c,g-i).

**Figure 2:**
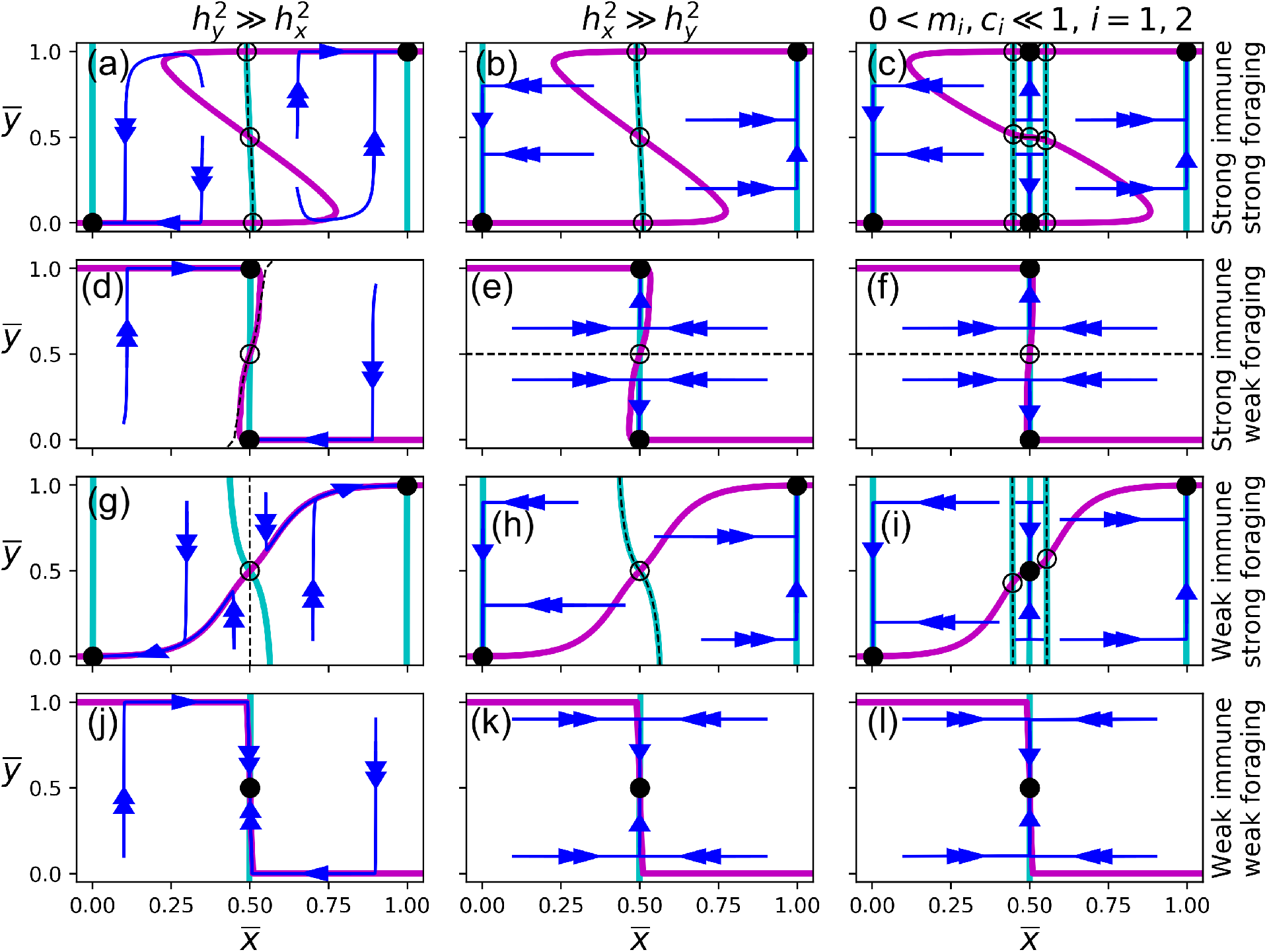
Nullclines, stable and unstable equilibria, separatrices, and evolutionary dynamics of (2). The cyan and pink curves denote the 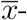- and 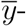nullclines, respectively. Filled-in and hollow circles indicate stable and unstable evolutionary equilbria, respectively. Blue lines are sample trajectories, and dashed lines indicate separatrices between stable equilibria (as well as the stable manifolds of the saddles). In (a), (d), (g), and (j), *σ*_*x,G*_ = 0.001 and *σ*_*y,G*_ = 0.25. In (b), (e), (h), and (k), *σ*_*x,G*_ = 0.25 and *σ*_*y,G*_ = 0.001. In (c), (f), (i), and (l), *m*_*i*_ = *c*_*i*_ = 0.1 for *i* = *L, B*. In (a)-(f) immune tradeoffs are strong (*τ*_*i*_ = 0.01), and in (g)-(l) immune tradeoffs are weak (*τ*_*i*_ = 1). In the (a)-(c) and (g)-(i) foraging tradeoffs are strong (*ζ*_*i*_ = 0.01) and in (d)-(f) and (j)-(l) foraging tradeoffs are weak (*ζ*_*i*_ = 1).

Figure 2 shows the evolutionary dynamics of (2) for a variety of scenarios. It also high-lights three major asymmetries of the foraging and immune traits in the context of three-timescale dynamics: (i) the relationship between trait tradeoff and equilibrium location, (ii) the relationship between initial and final evolutionary state, and (iii) the directionality of trait evolution.

Predators evolve generalist foraging strategies if the foraging tradeoff is weak, regardless of the immune tradeoff (Figure 2d-f,j-l). In contast, predators evolve generalist immune strategies if the immune tradeoff is weak, but only when the predator already has a generalist foraging strategy (Figure 2i-l). The immune tradeoff needs to be very weak (in relation to the foraging tradeoff) in order to have the same effect as the foraging tradeoff.

The final foraging state is generally determined by the initial foraging state, regardless of the strengths of trait tradeoffs. In contrast, the final immune state depends on both initial foraging and immune states, the strength of the trait tradeoffs, and ecological parameters such as parasite prevalence and lethality. Mathematically, the 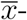nullclines in Figure (2) remain relatively vertical regardless of the strength of the foraging and immune tradeoffs, in contrast to the 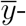nullclines, which are are never only horizontal. Consider for example a specialist predator (in both foraging and immune state) in an environment in which foraging and immune tradeoffs are strong (Figure 2a-c). The stabilizing selection at this evolutionary state is strong enough to withstand weakening immune tradeoffs, but not weakening foraging tradeoffs. In fact, if the foraging tradeoff becomes sufficiently weak, the predator will evolve a generalist foraging strategy, and an immune strategy dependent on the environment and the relative heritabilities of the two traits.

As a result of the extreme nature of the 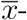nullclines, the foraging trait always evolves unidirectionally. On the other hand, because the 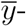nullclines are not strictly horizontal, the immune state may reverse its evolution when immune heritability is high relative to foraging heritability (Figure 2a,g,j). In these scenarios, the immune state evolves quickly toward a stable branch of the 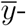nullcline, and then both traits evolve along the 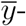nullcline toward a stable evolutionary equilibrium.

### Two-timescale dynamics

We used Latin hypercube sampling over a subset of parameter space to understand how the locations of the stable equilibria change as parameters vary (Figure 3). When both tradeoffs are strong (Figure 3a), equilibria congregate near evolutionary specialist states, while when both tradeoffs are weak (Figure 3d), the predator is more likely to evolve a generalist foraging and immune strategy. If foraging tradeoffs are weak and immune tradeoffs are strong (Figure 3b), predators will typically evolve a generalist foraging strategy and a specialist immune strategy. In contast, if immune tradeoffs are weak and foraging tradeoffs are strong (Figure 3c), then predators may evolve a generalist or specialist foraging strategy, and the immune strategy will evolve to correspond with the foraging state. We see the same asymmety as in Figure 2: generalist immune traits only evolve for generalist foragers, but generalist foraging traits may evolve regardless of immune state.

**Figure 3:**
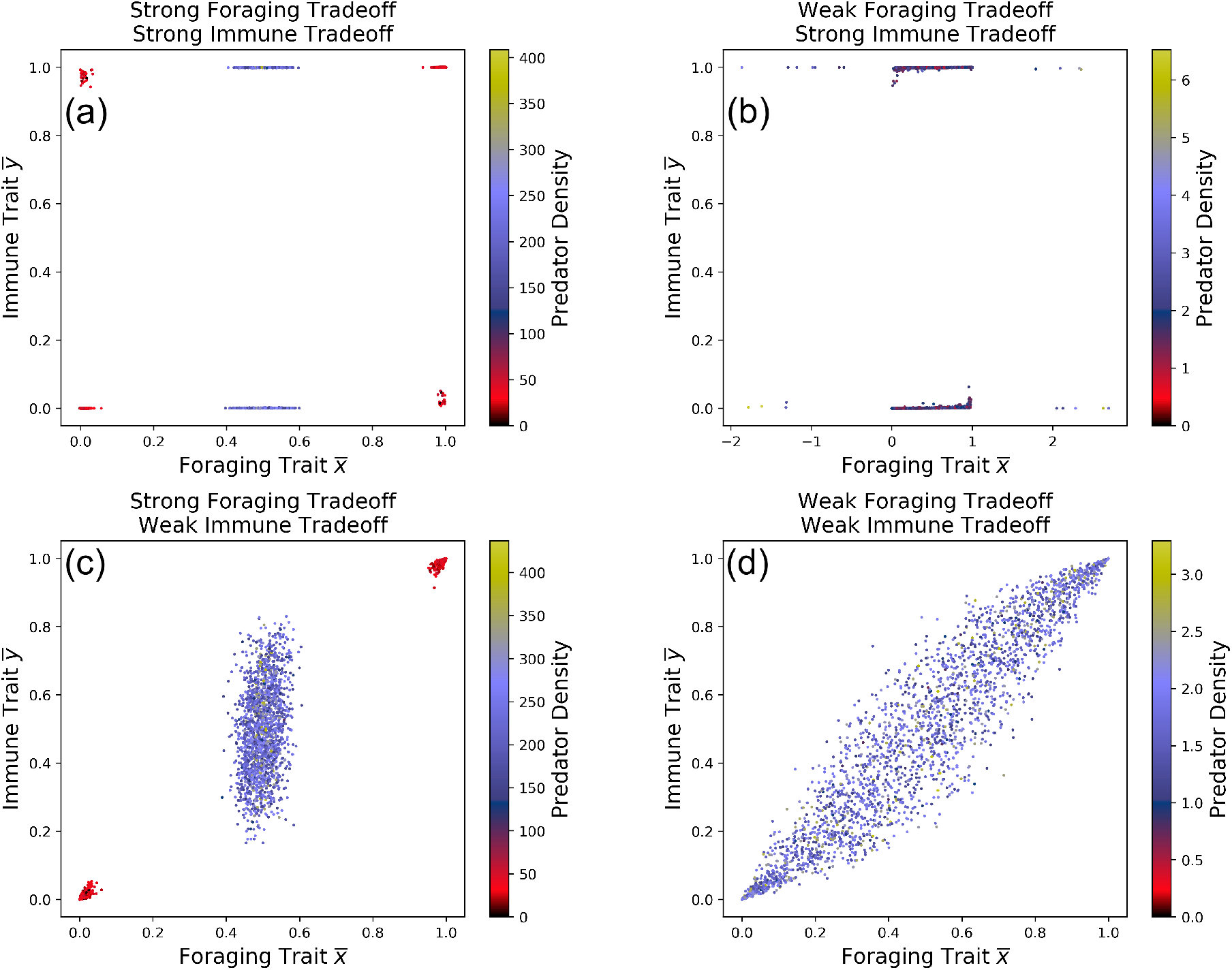
Locations of stable equilibria for a Latin Hypercube sample of parameter space. In (a) and (b) immune tradeoffs are strong (*τ*_*i*_ = 0.01), and in (c) and (d) immune tradeoffs are weak (*τ*_*i*_ = 1). In (a) and (c), foraging tradeoffs are strong (*ζ*_*i*_ = 0.01) and in (b) and (d) foraging tradeoffs are weak (*ζ*_*i*_ = 1). The color of each dot represents the density of the predator population at the evolutionary equilibrium. *K*_*i*_ ∈ [10, 1000], *α*_*i*_ ∈ [0.5, 0.9], *β*_*i*_ ∈ [0.9, 1], *γ*_*i*_ ∈ [0, 0.1], *b*_*i*_ ∈ [0.8, 1.2], *c*_*i*_, *m*_*i*_ ∈ [0.5, 1], *r*_1_ ∈ [0.5, 1.5], *d* ∈ [0.25, 0.55].

We also used the same Latin hypercube sample to understand what causes various correlations between prey intake, parasite exposure, parasite infection across predator populations (Figure 4). When immune tradeoffs are strong (Figure 4a,b), the relationship between prey intake and parasite infection does not stray far from the one-to-one line. In these scenarios, the proportion of a predator population’s diet consisting of some prey is roughly equal to the proportion of that predator’s parasite load consisting of the parasites from that prey. In addition, the majority of the variation in parasite infection is caused by the relationship between prey intake and parasite exposure, and not between exposure and infection. This means that any potential nonlinear pattern between diet and infection is not caused by immune evolution if immune tradeoffs are strong.

**Figure 4:**
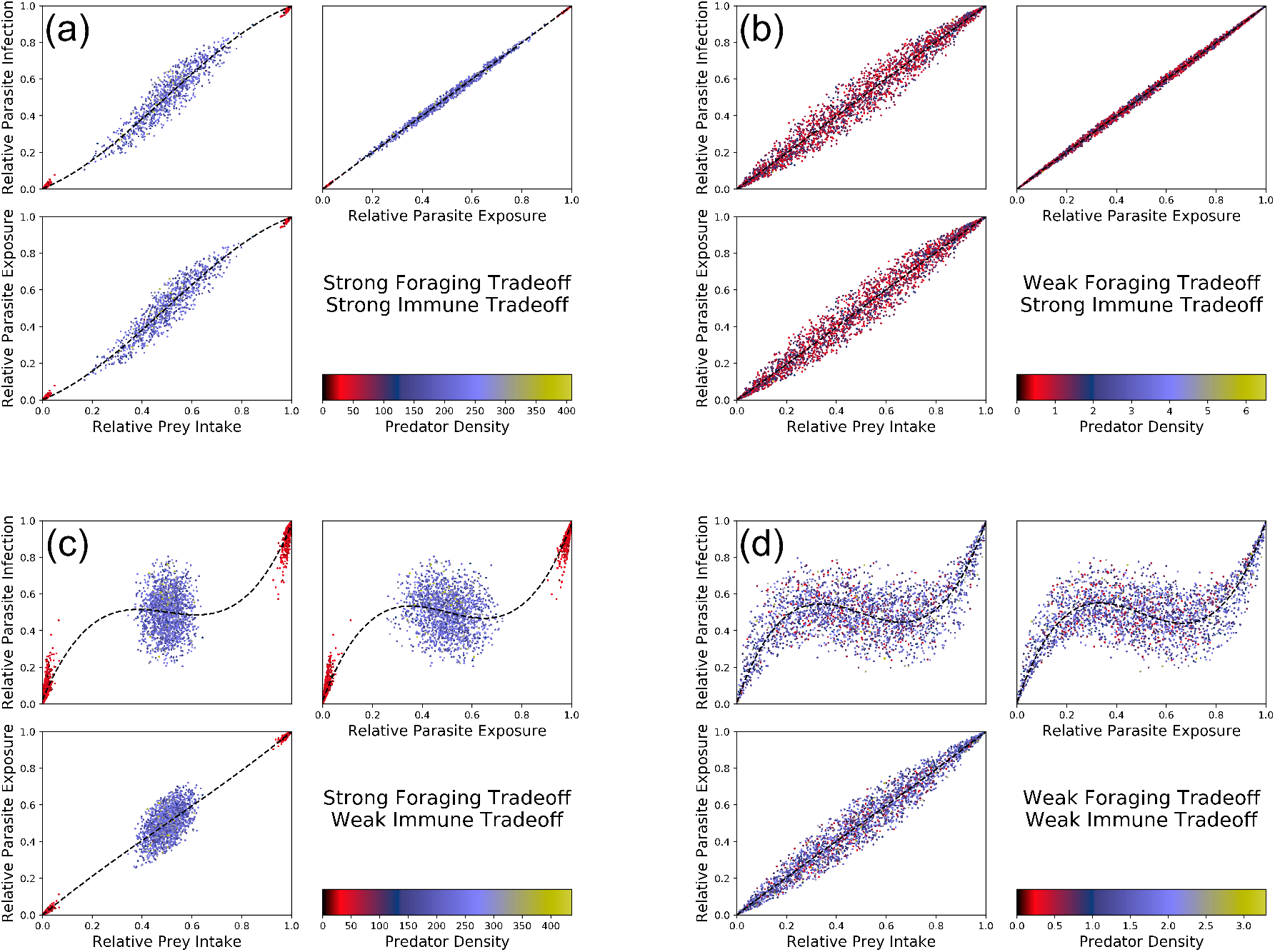
Prey intake, parasite exposure, and parasite infection over the same subset of parameter space given in Figure 3. The dots are colored as in Figure 3. The blue lines are splines of the data, included in order to better identify patterns between prey intake, parasite exposure, and parasite infection. In (a) and (b) immune tradeoffs are strong (*τ*_*i*_ = 0.01), and in (c) and (d) immune tradeoffs are weak (*τ*_*i*_ = 1). In (a) and (c), foraging tradeoffs are strong (*ζ*_*i*_ = 0.01) and in (b) and (d) foraging tradeoffs are weak (*ζ*_*i*_ = 1).

When immune tradeoffs are weak (Figure 4c,d), the relationship between prey intake and parasite infection differs greatly from the one-to-one line. In these scenarios, the proportion of a predator population’s diet consisting of some prey may not predict the proportion of that predator population’s parasite load consisting of the parasites from that prey. In contrast to when immune tradeoffs are strong, the majority of the variation in parasite infection *is* caused by the relationship between parasite exposure and parasite infection, indicating that any potential nonlinear pattern between diet and infection is caused by immune evolution if immune tradeoffs are weak.

### Non-equilibrium eco-evolutionary dynamics

When genetic heritability is high, eco-evolutionary feedbacks lead to greater dynamical complexity, including cyclical and chaotic dynamics. Schreiber et al. (2011) showed niche evolution can induce cycles and chaos in the absense of immune evolution, and we found something similar for immune evolution. Regardless of immune heritability, eco-evolutionary cycles only occur for sufficiently weak immune tradeoffs (Figure 5a). When immune heritability is low (Figure 5b), chaos occurs for very weak immune tradeoffs, while for higher immune heritability (Figure 5c,d), chaos occurs for intermediate and possible also very weak immune tradeoffs.

**Figure 5:**
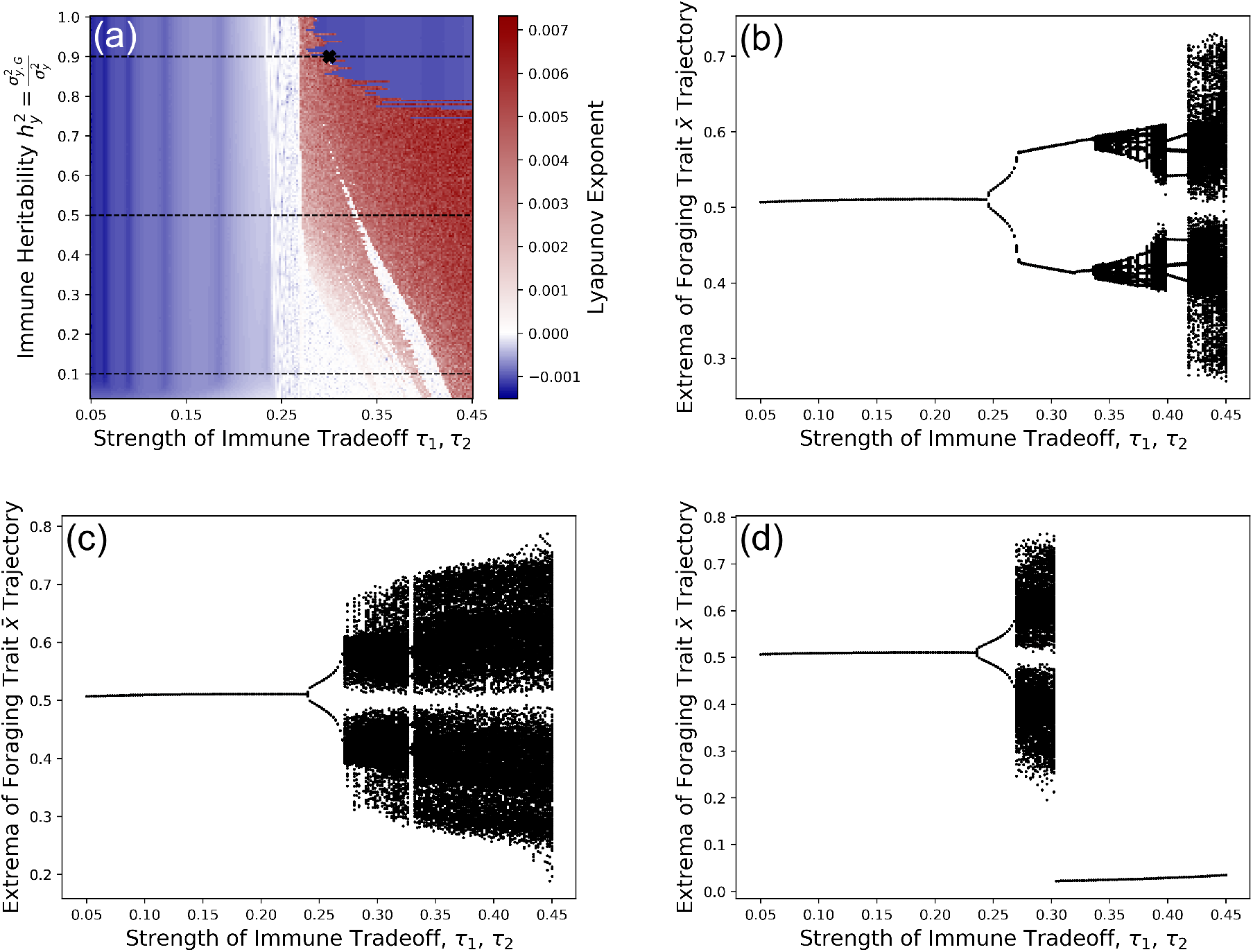
When immune tradeoffs are sufficiently weak, immune evolution can induce cyclic or chaotic eco-evolutionary dynamics. For weak immune tradeoffs, low heritability is destabilizing, but for intermediate immune tradeoffs, high heritability is destabilizing. (a) The blue regions denote stability (negative Lyapunov exponent) and the red regions denote chaos (positive Lyapunov exponent). (b) Local extrema of the niche trait 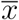 along 0.05 ≤ *τ*_1_ = *τ*_2_ ≤ 0.45, 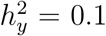. (c) Local extrema of the niche trait 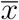 along 0.05 *≤ τ*_1_ = *τ*_2_ *≤* 0.45, 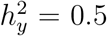. (d) Local extrema of the niche trait 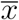 along 0.05 *≤ τ*_1_ = *τ*_2_ *≤* 0.45, 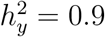.

A typical chaotic eco-evolutionary trajectory is displayed in Figure 6. These dynamics show a positive temporal correlation between foraging and immune traits (Figure 6c,d).

**Figure 6:**
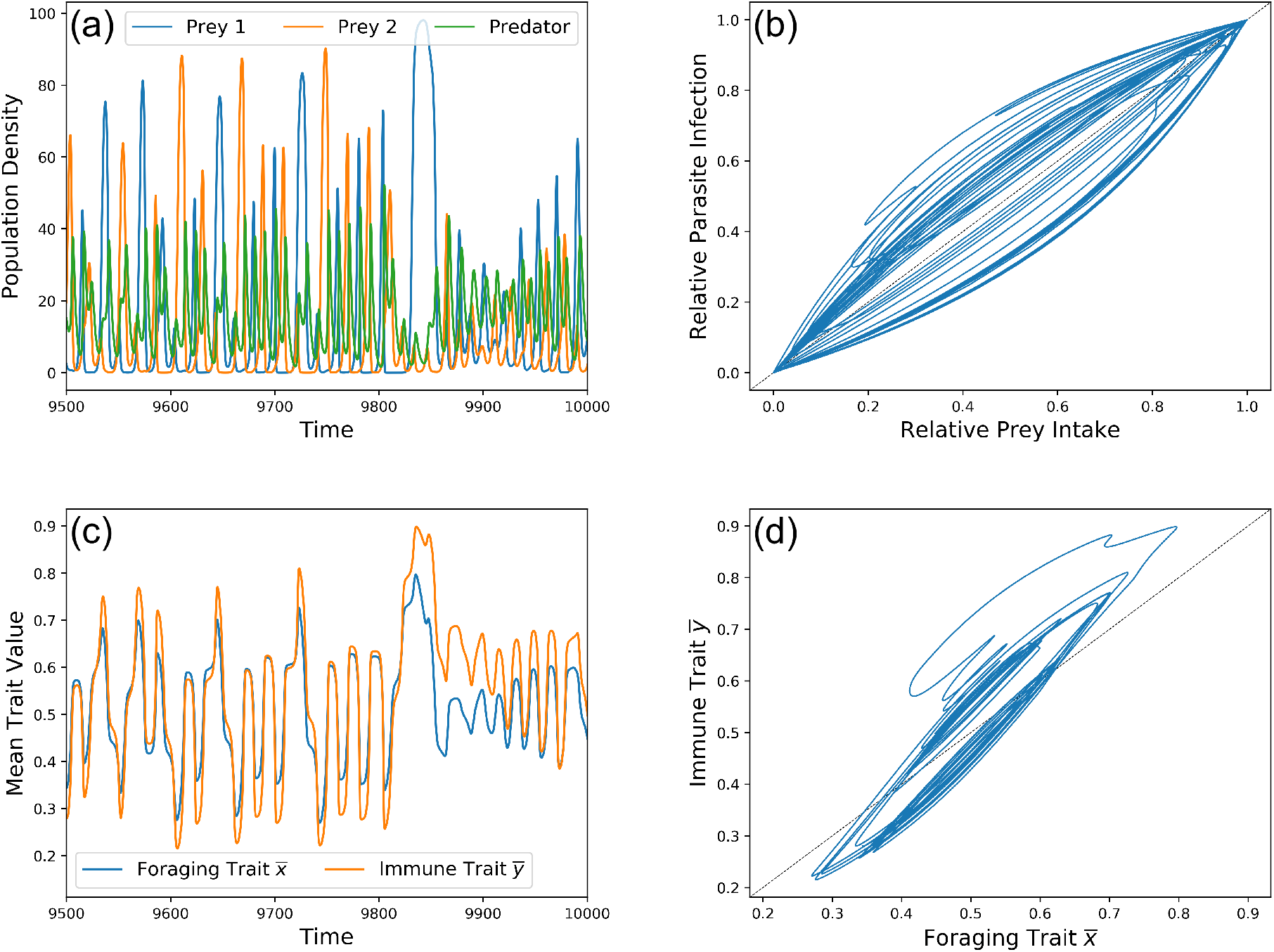
A chaotic eco-evolutionary trajectory. Parameters are equal to that of Figure 5 with *τ*_1_ = *τ*_2_ = 0.3 and 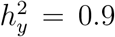 (bold x in Figure 5a). (a) predator and prey densities through time. (b) Predator relative prey intake vs. relative parasite infection. (c) predator foraging and immune traits through time. (d) predator foraging trait vs. immune trait.

When the predator foraging trait favors one prey type over the other, its intake almost entirely consists of that prey. The immune trait has higher heritability than that the foraging trait, which is why the immune trait evolves more extreme values than the foraging trait. Once the predator over-consumes a particular prey type and the other recovers, the foraging trait faces directional selection toward the recovering, although there is a lag in the actual intake of that prey. Although it is highly heritable, the immune trait does not favor the parasite of the recovering prey until the foraging trait is near its extreme value.

There is also a nonlinear correlation between diet and infection (Figure 6b). Because this correlation occurs within a single oscillating population, the intake-exposure relationship is one-to-one. Thus, any nonlinear correlation between diet and infection is entirely caused by the evolving immune trait 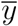.

## Discussion

In this study we addressed three questions: (a) how the evolution of a predator’s trophic niche affects evolution of immunity to trophically transmitted parasites, and vice-versa, (b) how these traits are correlated across populations and within a single population undergoing eco-evolutionary oscillations, and (c) how the dual evolution of niche and immunity can affect the correlation between intake of and infection by parasites among populations and within a single population undergoing eco-evolutionary oscillations.

To answer (a), we used mathematical techniques involving timescale separation to describe conditions under which niche evolution determines immune evolution and vice-versa. We found that even when the predator’s immune state evolves faster than it’s niche (due to high immune heritability and low ecomorphology heritability), immune evolution does not determine niche. On the other hand, niche evolution often determines the predator’s immune state, regardless of the relative speeds of evolution. Indeed, a fast-evolving immune state may reverse its evolution in response to a shifting niche, but the niche does not respond in kind to a shifting immunity.

The asymmetry between niche and immunity extends beyond the question of which trait determines the other. When niche tradeoffs are weak and immune tradeoffs are strong, the predator maintains a generalist foraging ecomorphology even though its immunity is specialized against one parasite or the other. This is true even if the parasites are abundant and detrimental to the predator’s fitness. On the other hand, when niche tradeoffs are strong and immune tradeoffs are weak, the predator evolves a specialist foraging ecomorphology along with a specialist immunity against the parasites it encounters. In short, there is little fitness benefit in maintaining an immunity to a parasite rarely encountered, but there is significant benefit to maintaining a morphology suitable to consume multiple prey even when they contain parasites that confer significant fitness drawbacks.

We also simulated eco-evolutionary dynamics on commensurate timescales to determine how an evolving immune trait can affect stability. While we did not calculate analytic conditions for stability or chaos, numerical results suggest that oscillations do not occur when immunological trdaeoffs are sufficiently strong. Theory predicts evolutionary destabilization occurs more commonly when there is a tradeoffs between capturing different prey phenotypes (Abrams 2000; Abrams and Matsuda 1997a,b), but we find that there is a limit to this effect; if tradeoffs are too strong, all evolutionary oscillations are suppressed.

To answer (b) and (c), we used Latin hypercube sampling to analyze how trophic and immune traits correlate, and how parasite intake and infection correlate, among populations. We found that there is no correlation between trophic and immune traits when immune tradeoffs are strong. In this scenario, predators will always evolve a specialized immunity regardless of their ecomorphology. When immune tradeoffs are weak, predators evolve an immunity to suit their niche. In this scenario, the strength of foraging tradeoffs determine the correlation; if foraging tradeoffs are strong, then predators evolve to only a few morphological states, whereas if foraging tradeoffs are weak, then predators may evolve anywhere along the ecomorphology spectrum.

Because predators only evolve immunity to suit their niche if immune tradeoffs are weak, negative correlations between parasite intake and infection are only possible in this scenario. The negative correlation between intake and infection is more pronounced when there is more morphological diversity across populations, and thus is most likely to be observed if foraging tradeoffs are also weak. These results, when compared to the empirical data of Stutz et al. (2014), suggest that evolutionary trade-offs in stickleback niche (benthic or limnetic prey) and immune traits (benthic or limnetic parasite) are likely to be weak. This aligns with other studies which have shown that many stickleback populations have evolved a generalist morphology when both limnetic and benthic prey are present (Lavin and McPhail 1985; Matthews et al. 2010; Schluter and McPhail 1992; Snowberg et al. 2015).

We also found a nonlinear correlation between intake and infection within a single population oscillating in time. Because of the assumption that parasite abundance stays constant, this correlation is caused entirely by the evolution of immunity. Our simulations did not produce a negative correlation between intake and infection within a single oscillating population. However, the nonlinear correlation suggests that the negative correlation between diet and infection across populations observed by Stutz et al. (2014) may have resulted from oscillating populations in similar habitats rather than equilibriated populations in different habitats. However, because ecological variation among lakes is correlated with lake size (larger lakes containing more limnetic-feeding populations), this is unlikely to be the case.

It is well known that stickleback face biomechanical trade-offs that limit their ability to capture both benthic and limnetic prey (Robinson 2000). In contrast, it is not known whether stickleback immunity face comparable immunological trade-offs. That is, does immunity to benthic-derived parasites (e.g., nematodes) also confer protection to limnetic-derived parasites (e.g., Schistocephalus cestodes), or inhibit immunity to cestodes? In general, evidence suggests that different parasites are detected by different host MHC IIb alleles (Stutz and Bolnick 2017), suggesting a possible trade-off. For certain kinds of parasites this trade-off is well documented, such as the mutual inhibition of Th1 and Th2 adaptive immune responses that, respectively, target bacterial and helminth infections. We chose to model stickleback immunity on a single bidirectional axis, with different optimal values for immunity against limnetic and benthic parasites. This choice comes with the implicit assumption that there is a limited amount of energy allocated toward immunity, but the vertebrate immune system is complex and highly multivariate. An alternative modeling framework we considered is one unidirectional axis describing immunity for limnetic parasites, and a separate unidirectional axis for immunity to benthic parasites. Unlike our modeling framework, trade-offs would not be between immunity to either limnetic or benthic parasites, but rather between immunity to a parasite species and some other unrelated metabolic rate (i.e. death rate or conversion efficiency).

We did not choose this framework for two reasons. First, this choice would push the evolutionary dimensionality of our model from two to three, and the total dimensionality from five to six, thus increasing the complexity of analysis and decreasing mathematical tractibility. Also, we expect that negative correlations between intake and infection are more likely to be seen in this more complex modeling framework. This is because there is no trade-off between immune traits, and thus slight evolutionary adjustments in immunity to one parasite can be made without affecting immunity to the other. Thus, in this framework, even when the trade-off between immunity and a metabolic rate is strong, populations can optimize immunity along both axes, allowing for a kind of immunity generalism. When populations are able to more finely tune their immunity, negative correlations between intake and infection are more likely. We were able to produce negative correlations between intake and infection in a far more conservative framework, less likely to produce this correlation, which is strong evidence in favor of the hypothesis of Stutz et al. (2014) and suggests that the cause of the negative correlation between intake and infection is indeed the evolution of immune traits in conjunction with trophic niche traits.

This study was motivated in part by the specific relationship between stickleback and their infected prey. However, trophically transmitted parasites are very common in nature (Combes 2001). This study helps shed light on how food web dynamics are affected by the presence of diet-derived parasites, and is a contribution to the growing body of theory regarding eco-evo dynamics in a multispecies context (Abrams and Matsuda 1997b; Abrams 2006; Cortez and Weitz 2014; Cortez and Patel 2017; Fleischer et al. 2018; Klauschies et al. 2016; Patel and Bürger 2019; Patel and Schreiber 2015, 2018; Saloniemi 1993; Schreiber et al. 2011; Schreiber and Patel 2015; Tien and Ellner 2012; Vasseur and Fox 2011, and others). In particular, predator traits evolve in response to the presence and danger of parasites in prey, which results in a shift in predator exposure and susceptibility to parasites. These eco-evo feedbacks can cause chaos or other oscillatory behavior, or cause alternative stable ecological and evolutionary states.

Conversely, we need improve our understanding of how food web dynamics play a role in the dynamics of trophically transmitted parasites, and future theoretical studies should incorporate the dynamics of parasites along with the dynamics of the community in which they reside. Prosnier et al. (2020) used an epidemiological framework to examine the effect of prey infection on predator diet. They also examined the effect of predator diet evolution on coexistence using an adaptive dynamics evolutionary framework and showed that this type of evolution generally promotes coexistence among a predator and an infected and uninfected prey. Like in this study, reductions in prey density correspond to lower consumption rates by the predator, which ultimately favors prey persistence.

Prosnier et al. (2020) modeled parasite transmission as horizontal between prey. This is not the case for stickleback prey, which become infected by parasites through consumption (Barber and Scharsack 2009). We therefore chose not to include explicit parasite dynamics or the epidemiological dynamics of the prey, and instead assumed that the proportion of prey which are infected stays constant. Common predators of stickleback are piscivorous birds which freely move between many lakes or ponds. Parasites lay eggs in the gut of a bird and these eggs are deposited into lakes when these birds defacate above water. This suggests that a significant proportion of the parasite load in prey results from regional recruitment rather than local population reproduction. That is, birds which consume infected stickleback from one lake or pond often defacate into another, which keeps the parasite load in each lake or pond relatively constant.

Although it may be that immunity and ecomorphology are genetically linked in some way, we chose to model the two traits as genetically independent. This choice improves mathematical tractibility, as well as provides an example of a system in which the evolution of two traits drive each other, not because of genetic linkage, but rather based solely on interdependent selection pressures. Future studies should explore how correlated selection pressures can enhance, or reverse, the effects of genetic linkage of two traits.

Finally, experiments are needed to validate our model, including measurements of relevant parameters and tests of our assumptions. We know that stickleback individuals vary in their propensity to consume benthic versus limnetic resources (Bolnick and Lau 2008; Bolnick et al. 2014; Matthews et al. 2010; Robinson 2000; Snowberg et al. 2015). However, the precise nature and strength of the biomecanical (and perhaps cognitive) trade-offs remain poorly understood (Robinson 2000; Schmid et al. 2019). Likewise, we know that stickleback genotypes differ in their resistance to various parasites (Eizaguirre et al. 2012; Kalbe and Kurtz 2005; MacColl 2009; MacColl and Chapman 2010; Nagar and MacColl 2016; Stutz and Bolnick 2017; Weber et al. 2017a,b, among many others). But, we know little about trade-offs (or synergy) between resistance to different parasites. For that matter, parasites can manipulate host immunity in ways that benefits or harms co-infecting parasites (e.g., Ezenwa et al. 2010). We therefore need to bring together biomechanical studies of foraging trade-offs, with mechanistic immunological studies of resistance trade-offs. In addition, we lack sufficient information about the relative virulence of different parasites acquired through alternative prey, and future theoretical studies should include the effects and evolution of all three host strategies: avoidance, resistance, and tolerance.

## References

[1] P. A. Abrams. “The Evolution of Predator-Prey Interactions: Theory and Evidence”. In: Annual Review of Ecology and Systematics 31 (2000), pp. 79–105.

[2] P. A. Abrams and H. Matsuda. “Fitness minimization and dynamic instability as a consequence of predator–prey coevolution”. In: Evolutionary Ecology 11.1 (Jan. 1997), pp. 1–20.

[3] P. A. Abrams and H. Matsuda. “Prey Adaptation as a Cause of Predator-Prey Cycles”. In: Society for the Study of Evolution 51 (1997), pp. 1742–1750.

[4] P. A. Abrams. “Adaptive change in the resource-exploitation traits of a generalist consumer: the evolution and coexistence of generalists and specialists”. In: Evolution 60.3 (Mar. 2006), pp. 427–439.

[5] T. K. Anderson and M. V. K. Sukhdeo. “Host Centrality in Food Web Networks Determines Parasite Diversity”. In: PLOS ONE 6.10 (Oct. 2011), pp. 1–9.

[6] I. Barber and J. P. Scharsack. “The three-spined stickleback-Schistocephalus solidus system: an experimental model for investigating host-parasite interactions in fish”. In: Parasitology 137(3) (2009), pp. 411–424.

[7] L. Becks, S. P. Ellner, L. E. Jones, and N. G. Hairston Jr. “Reduction of adaptive genetic diversity radically alters eco-evolutionary community dynamics”. In: Ecology Letters 13.8 (2010), pp. 989–997.

[8] D. I. Bolnick and O. L. Lau. “Predictable Patterns of Disruptive Selection in Stickleback in Postglacial Lakes.” In: The American Naturalist 172.1 (2008). PMID: 18452402, pp. 1–11.

[9] D. I. Bolnick, L. K. Snowberg, P. E. Hirsch, C. L. Lauber, R. Knight, J. G. Caporaso, and R. Svanbäck. “Individuals’ diet diversity influences gut microbial diversity in two freshwater fish (threespine stickleback and Eurasian perch)”. In: Ecology Letters 17.8 (May 2014). Ed. by D. Post, pp. 979–987.

[10] C. Combes. Parasitism: The Ecology and Evolution of Intimate Interactions. University of Chicago Press, 2001.

[11] M. H. Cortez and J. S. Weitz. “Coevolution can reverse predator–prey cycles”. In: Proceedings of the National Academy of Sciences 111.20 (2014), pp. 7486–7491.

[12] M. H. Cortez and S. Patel. “The Effects of Predator Evolution and Genetic Variation on Predator–Prey Population-Level Dynamics”. In: Bulletin of Mathematical Biology 79.7 (June 2017), pp. 1510–1538.

[13] C. Eizaguirre, T. L. Lenz, M. Kalbe, and M. Milinski. “Divergent selection on locally adapted major histocompatibility complex immune genes experimentally proven in the field”. In: Ecology Letters 15.7 (May 2012). Ed. by G. Sorci, pp. 723–731.

[14] J. A. Estes, M. L. Riedman, M. M. Staedler, M. T. Tinker, and B. E. Lyon. “Individual variation in prey selection by sea otters: patterns, causes and implications”. In: Journal of Animal Ecology 72.1 (Jan. 2003), pp. 144–155.

[15] V. O. Ezenwa, R. S. Etienne, G. Luikart, A. Beja-Pereira, and A. E. Jolles. “Hidden Consequences of Living in a Wormy World: Nematode-Induced Immune Suppression Facilitates Tuberculosis Invasion in African Buffalo”. In: The American Naturalist 176.5 (Nov. 2010), pp. 613–624.

[16] S. R. Fleischer, C. P. terHorst, and J. Li. “Pick your trade-offs wisely: Predator-prey eco-evo dynamics are qualitatively different under different trade-offs”. In: Journal of Theoretical Biology 456 (Nov. 2018), pp. 201–212.

[17] A. Hayward, M. Tsuboi, C. Owusu, A. Kotrschal, S. D. Buechel, J. Zidar, C. K. Cornwallis, H. Løvlie, and N. Kolm. “Evolutionary associations between host traits and parasite load: insights from Lake Tanganyika cichlids”. In: Journal of Evolutionary Biology 30.6 (Mar. 2017), pp. 1056–1067.

[18] R. Iritani and T. Sato. “Host-Manipulation by Trophically Transmitted Parasites: The Switcher-Paradigm”. In: Trends in Parasitology 34.11 (Nov. 2018), pp. 934–944.

[19] C. K. Johnson, M. T. Tinker, J. A. Estes, P. A. Conrad, M. Staedler, M. A. Miller, D. A. Jessup, and J. A. K. Mazet. “Prey choice and habitat use drive sea otter pathogen exposure in a resource-limited coastal system”. In: Proceedings of the National Academy of Sciences 106.7 (Jan. 2009), pp. 2242–2247.

[20] E. Jones, T. Oliphant, P. Peterson, et al. SciPy: Open source scientific tools for Python. [Online; accessed 20 May 2019]. 2001–.

[21] M. Kalbe and J. Kurtz. “Local differences in immunocompetence reflect resistance of sticklebacks against the eye fluke Diplostomum pseudospathaceum”. In: Parasitology 132.01 (Sept. 2005), pp. 105–116.

[22] T. Klauschies, D. A. Vasseur, and U. Gaedke. “Trait adaptation promotes species coexistence in diverse predator and prey communities”. In: Ecology and Evolution 6.12 (2016), pp. 4141–4159.

[23] K. D. Lafferty, A. P. Dobson, and A. M. Kuris. “Parasites dominate food web links”. In: Proceedings of the National Academy of Sciences 103.30 (2006), pp. 11211–11216.

[24] R. Lande. “Natural Selection and Random Genetic Drift in Phentypic Evolution”. In: Society for the Study of Evolution 30(2) (1976), pp. 314–334.

[25] P. Lavin and J. D. McPhail. “Adaptive divergence of trophic phenotype among fresh-water populations of the threespine stickleback (Gasterosteus aculeatus)”. In: Canadian Journal of Fisheries and Aquatic Sciences 43 (1986), pp. 2455–2463.

[26] P. Lavin and J. D. McPhail. “The evolution of freshwater diversity in the threespine stickleback (Gasterosteus aculeatus): site-specific differentiation of trophic morphology”. In: Canadian Journal of Zoology 83 (1985), pp. 2632–2638.

[27] A. D. C. MacColl. “Parasite burdens differ between sympatric three-spined stickleback species”. In: Ecography 32.1 (Apr. 2009), pp. 153–160.

[28] A. D. C. MacColl and S. M. Chapman. “Parasites can cause selection against migrants following dispersal between environments”. In: Functional Ecology 24.4 (Feb. 2010), pp. 847–856.

[29] B. Matthews, K. B. Marchinko, D. I. Bolnick, and A. Mazumder. “Specialization of trophic position and habitat use by sticklebacks in an adaptive radiation”. In: Ecology 91 (2010), pp. 1025–1034.

[30] A. E. Nagar and A. D. C. MacColl. “Parasites contribute to ecologically dependent postmating isolation in the adaptive radiation of three-spined stickleback”. In: Proceedings of the Royal Society B: Biological Sciences 283.1836 (Aug. 2016), p. 20160691.

[31] S. Patel and R. Bürger. “Eco-evolutionary feedbacks between prey densities and linkage disequilibrium in the predator maintain diversity”. In: Evolution 73.8 (June 2019), pp. 1533–1548.

[32] S. Patel and S. J. Schreiber. “Evolutionarily Driven Shifts in Communities with Intraguild Predation”. In: The American Naturalist 186.5 (Nov. 2015), E98–E110.

[33] S. Patel and S. J. Schreiber. “Robust permanence for ecological equations with internal and external feedbacks”. In: Journal of Mathematical Biology 77.1 (2018), pp. 79–105.

[34] L. Prosnier, V. Médoc, and N. Loeuille. “Evolution of predator foraging in response to prey infection favors species coexistence”. In: bioRxiv (2020).

[35] B. Robinson. “Trade Offs in Habitat-Specific Foraging Efficiency and the Nascent Adaptive Divergence of Sticklebacks in Lakes”. In: Behaviour 137.7-8 (2000), pp. 865–888.

[36] A. Rogawa, S. Ogata, and A. Mougi. “Parasite transmission between trophic levels stabilizes predator–prey interaction”. In: Scientific Reports 8.1 (Aug. 2018).

[37] I. Saloniemi. “A Coevolutionary Predator-Prey Model with Quantitative Characters.” In: The American Naturalist 141 (1993), pp. 880–896.

[38] D. Schluter and J. D. McPhail. “Ecological character displacement and speciation in sticklebacks”. In: The American Naturalist 140 (1992), pp. 85–108.

[39] D. W. Schmid, M. D. McGee, R. J. Best, O. Seehausen, and B. Matthews. “Rapid Divergence of Predator Functional Traits Affects Prey Composition in Aquatic Communities”. In: The American Naturalist 193.3 (Mar. 2019), pp. 331–345.

[40] S. J. Schreiber, R. Bürger, and D. I. Bolnick. “The community effects of phenotypic and genetic variation within a predator population”. In: Ecology 92.8 (2011), pp. 1582–1593.

[41] S. J. Schreiber and S. Patel. “Evolutionarily induced alternative states and coexistence in systems with apparent competition”. In: Natural Resource Modeling 28.4 (2015), pp. 475–496.

[42] F. A. Sibbing, L. A. J. Nagelkerke, R. J. M. Stet, and J. W. M. Osse. In: Aquatic Ecology 32.3 (1998), pp. 217–227.

[43] L. K. Snowberg, K. M. Hendrix, and D. I. Bolnick. “Covarying variances: more morphologically variable populations also exhibit more diet variation”. In: Oecologia 178(1) (2015), pp. 89–101.

[44] J. C. Sprott. Chaos and time-series analysis. Vol. 69. Citeseer, 2003.

[45] M. Stein. “Large Sample Properties of Simulations Using Latin Hypercube Sampling”. In: Technometrics 29.2 (1987), pp. 143–151.

[46] W. E. Stutz and D. I. Bolnick. “Natural selection on MHC IIβ in parapatric lake and stream stickleback: Balancing, divergent, both or neither?” In: Molecular Ecology 26.18 (May 2017), pp. 4772–4786.

[47] W. E. Stutz, O. L. Lau, and D. I. Bolnick. “Contrasting Patterns of Phenotype-Dependent Parasitism within and among Populations of Threespine Stickleback.” In: The American Naturalist 183.6 (2014), pp. 810–825.

[48] M. V. K. Sukhdeo. “Where are the parasites in food webs?” In: Parasites & Vectors 5.1 (2012), p. 239.

[49] R. J. Tien and S. P. Ellner. “Variable cost of prey defense and coevolution in predator–prey systems”. In: Ecological Monographs 82.4 (2012), pp. 491–504.

[50] M. T., Tinker, B. G., and J. A. Estes. “Food limitation leads to behavioral diversification and dietary specialization in sea otters”. In: Proceedings of the National Academy of Sciences 105.2 (Jan. 2008), pp. 560–565.

[51] D. A. Vasseur and J. W. Fox. “Adaptive Dynamics of Competition for Nutritionally Complementary Resources: Character Convergence, Displacement, and Parallelism.” In: The American Naturalist 178.4 (2011). PMID: 21956028, pp. 501–514.

[52] E. van Velzen and U. Gaedke. “Disentangling eco-evolutionary dynamics of predator-prey coevolution: the case of antiphase cycles”. In: Scientific Reports 7.1 (2017), p. 17125.

[53] J. N. Weber, M. Kalbe, K. C. Shim, N. I. Erin, N. C. Steinel, L. Ma, and D. I. Bolnick. “Resist Globally, Infect Locally: A Transcontinental Test of Adaptation by Stickleback and Their Tapeworm Parasite”. In: The American Naturalist 189.1 (Jan. 2017), pp. 43–57.

[54] J. N. Weber, N. C. Steinel, K. C. Shim, and D. I. Bolnick. “Recent evolution of extreme cestode growth suppression by a vertebrate host”. In: Proceedings of the National Academy of Sciences 114.25 (June 2017), pp. 6575–6580.

[55] C. L. Wood and P. T. J. Johnson. “A world without parasites: exploring the hidden ecology of infection”. In: Frontiers in Ecology and the Environment 13.8 (2015), pp. 425–434.

[56] T. Yoshida, L. E. Jones, S. P. Ellner, G. F. Fussmann, and N. G. Hairson Jr. “Rapid evolution drives ecological dynamics in a predator–prey system”. In: Nature 424 (2003), pp. 303–306.

