## Appendix A for "Sick of Eating: eco-evo-immuno dynamics of predators and their trophically acquired parasites"

### Sick of Eating: eco-evo-immuno dynamics of predators and their trophically acquired parasites
### Appendix A: Derivation of average prey and predator fitness

Samuel R. Fleischer*[1], Daniel I. Bolnick[2], and Sebastian J. Schreiber[3]

[1]*Graduate Group in Applied Mathematics, University of California, Davis*

[2]*Department of Ecology and Evolutionary Biology, University of Connecticut*

[3]*Department of Evolution and Ecology, University of California, Davis*

May 26, 2020

Recall the probability distributions $p_x(x, \overline{x})$ and $p_y(y, \overline{y})$:

$$p_x(x, \overline{x}) = \frac{1}{\sqrt{2\pi\sigma_x^2}} \exp\left[-\frac{(x-\overline{x})^2}{2\sigma_x^2}\right] \qquad \text{and} \qquad p_y(y, \overline{y}) = \frac{1}{\sqrt{2\pi\sigma_y^2}} \exp\left[-\frac{(y-\overline{y})^2}{2\sigma_y^2}\right] \tag{A1}$$

Given $W(x, y) = \sum_{i=1}^{2} \Big[(b_i - c_i m_i S_i(y)) a_i(x) N_i\Big] - d$, then through the linearity of sums and integrals, we have

$$\overline{W}(\overline{x}, \overline{y}) = \int_{\mathbb{R}\times\mathbb{R}} W(x, y) p_x(x, \overline{x}) p_y(y, \overline{y}) \mathrm{d}x\mathrm{d}y = \sum_{i=1}^{2} \Big[(b_i - c_i m_i \overline{S}_i(\overline{y})) \overline{a}_i(\overline{x}) N_i\Big] - d \tag{A2}$$

where

$$\overline{a}_i(\overline{x}) = \int_{\mathbb{R}} a_i(x) p_x(x, \overline{x}) \mathrm{d}x \qquad \text{and} \qquad \overline{S}_i(\overline{y}) = \int_{\mathbb{R}} S_i(y) p_y(y, \overline{y}) \mathrm{d}y \tag{A3}$$

are the average attack rate of the predator on prey $i$ and the susceptibility rate of the predator to infection by the parasite in prey $i$, respectively. Similary, given $Y_i(x) = r_i\left(1 - \dfrac{N_i}{K_i}\right) - a_i(x)P$ for $i = 1, 2$, we have

$$\overline{Y}_i(\overline{x}) = \int_{\mathbb{R}} Y_i(x) p_x(x, \overline{x}) \mathrm{d}x = r_i\left(1 - \frac{N_i}{K_i}\right) - \overline{a}_i(\overline{x})P. \tag{A4}$$

Finally, since

$$a_i(x) = \alpha_i \exp\left[-\frac{(x-\theta_i)^2}{2\zeta_i^2}\right] \qquad \text{and} \qquad S_i(y) = \beta_i - (\beta_i - \gamma_i)\exp\left[-\frac{(y-\phi_i)^2}{2\tau_i^2}\right], \tag{A5}$$

we have

$$\overline{a}_i(\overline{x}) = \int_{\mathbb{R}} \alpha_i \exp\left[-\frac{(x-\theta_i)^2}{2\zeta_i^2}\right] \frac{1}{\sqrt{2\pi\sigma_x^2}} \exp\left[-\frac{(x-\overline{x})^2}{2\sigma_x^2}\right] \mathrm{d}x \tag{A6}$$

$$= \frac{\alpha_i}{\sqrt{2\pi\sigma_x^2}} \int_{\mathbb{R}} \exp\left[-\frac{(x-\theta_i)^2}{2\zeta_i^2} - \frac{(x-\overline{x})^2}{2\sigma_x^2}\right] \mathrm{d}x \tag{A7}$$

$$= \frac{\alpha_i}{\sqrt{2\pi\sigma_x^2}} \int_{\mathbb{R}} \exp\Big[-\frac{1}{2\zeta_i^2\sigma_x^2}\big((\sigma_x^2 + \zeta_i^2)x^2 - (2\theta_i\sigma_x^2 + 2\overline{x}\zeta_i^2)x + (\theta_i^2\sigma_x^2 + \overline{x}^2\zeta_i^2)\big)\Big] \mathrm{d}x \tag{A8}$$

$$= \frac{\alpha_i}{\sqrt{2\pi\sigma_x^2}} \frac{\sqrt{\pi}}{\sqrt{A}} \exp\left[\frac{B^2}{4A} - C\right] \tag{A9}$$

where $A = \dfrac{\sigma_x^2 + \zeta_i^2}{2\zeta_i^2\sigma_x^2}$, $B = \dfrac{2\theta_i\sigma_x^2 + 2\overline{x}\zeta_i^2}{2\zeta_i^2\sigma_x^2}$, and $C = \dfrac{\theta_i^2\sigma_x^2 + \overline{x}^2\zeta_i^2}{2\zeta_i^2\sigma_x^2}$. So,

$$\frac{B^2}{4A} - C = -\frac{(\overline{x} - \theta_i)^2}{2(\sigma_x^2 + \zeta_i^2)} \tag{A10}$$

and thus

$$\overline{a}_i(\overline{x}) = \frac{\alpha_i \zeta_i}{\sqrt{\sigma_x^2 + \zeta_i^2}} \exp\left[-\frac{(\overline{x} - \theta_i)^2}{2(\sigma_x^2 + \zeta_i^2)}\right]. \tag{A11}$$

Also,

$$\overline{S}_i(\overline{y}) = \int_{\mathbb{R}} \left(\beta_i - (\beta_i - \gamma_i) \exp\left[-\frac{y - \phi_i}{2\tau_i^2}\right]\right) \frac{1}{\sqrt{2\pi\sigma_y^2}} \exp\left[-\frac{(y - \overline{y})^2}{2\sigma_y^2}\right] \mathrm{d}y \tag{A12}$$

$$= \beta_i \int_{\mathbb{R}} \frac{1}{\sqrt{2\pi\sigma_y^2}} \exp\left[-\frac{(y - \overline{y})^2}{2\sigma_y^2}\right] \mathrm{d}y - \frac{\beta_i - \gamma_i}{\sqrt{2\pi\sigma_y^2}} \int_{\mathbb{R}} \exp\left[-\frac{(y - \phi_i)^2}{2\tau_i^2} - \frac{(y - \overline{y})^2}{2\sigma_y^2}\right] \mathrm{d}y \tag{A13}$$

Through an identical calculation as for $\overline{a}_i(\overline{x})$, we have

$$\overline{S}_i(\overline{y}) = \beta_i - \frac{(\beta_i - \gamma_i)\tau_i}{\sqrt{\sigma_y^2 + \tau_i^2}} \exp\left[-\frac{(\overline{y} - \phi_i)^2}{2\left(\sigma_y^2 + \tau_i^2\right)}\right]. \tag{A14}$$
