## Appendix B for "Sick of Eating: eco-evo-immuno dynamics of predators and their trophically acquired parasites"

### Appendix B: Model parameters

Samuel R. Fleischer*[1], Daniel I. Bolnick[2], and Sebastian J. Schreiber[3]

[1]*Graduate Group in Applied Mathematics, University of California, Davis*

[2]*Department of Ecology and Evolutionary Biology, University of Connecticut*

[3]*Department of Evolution and Ecology, University of California, Davis*

May 26, 2020

| Parameter | Baseline Value |
|---|---|
| $K_1$, $K_2$ | 100 |
| $\alpha_1$, $\alpha_2$ | 0.7 |
| $\beta_1$, $\beta_2$ | 0.95 |
| $\gamma_1$, $\gamma_2$ | 0.05 |
| $\sigma_x$, $\sigma_y$ | 0.25 |
| $\sigma_{x,G}$, $\sigma_{y,G}$ | 0.1 |
| $\tau_1$, $\tau_2$ | 0.1 |
| $\zeta_1$, $\zeta_2$ | 0.1 |
| $\theta_1$, $\phi_1$ | 0 |
| $\theta_2$, $\phi_2$ | 1 |
| $b_1$, $b_2$ | 1 |
| $c_1$, $c_2$ | 0.9 |
| $m_1$, $m_2$ | 0.9 |
| $r_1$, $r_2$ | 1 |
| $d$ | 0.4 |

Table B1: Baseline parameter values.

| Figure(s) | Parameter | Value |
|---|---|---|
| Figure 2a-c,g-i | $\zeta_1$, $\zeta_2$ | 0.01 |
| Figure 2d-f,j-l | $\zeta_1$, $\zeta_2$ | 1 |
| Figure 2a-f | $\tau_1$, $\tau_2$ | 0.01 |
| Figure 2g-l | $\tau_1$, $\tau_2$ | 1 |
| Figure 2a,d,g,j | $\sigma_{x,G}$ | 0.005 |
| Figure 2a,d,g,j | $\sigma_{y,G}$ | 0.25 |
| Figure 2b,e,h,k | $\sigma_{x,G}$ | 0.25 |
| Figure 2b,e,h,k | $\sigma_{y,G}$ | 0.005 |
| Figure 2c,f,i,l | $c_1$, $c_2$, $m_1$, $m_2$ | 0.1 |

Table B2: Figure 2 parameters. All parameters not given here are given in Table B1.

| Figure(s) | Parameter | Value |
|---|---|---|
| Figures 3a-d and 4a-d | $K_1$, $K_2$ | $[10, 1000]$ |
| Figures 3a-d and 4a-d | $\alpha_1$, $\alpha_2$ | $[0.5, 0.9]$ |
| Figures 3a-d and 4a-d | $\beta_1$, $\beta_2$ | $[0.9, 1]$ |
| Figures 3a-d and 4a-d | $\gamma_1$, $\gamma_2$ | $[0, 0.1]$ |
| Figures 3a-d and 4a-d | $\sigma_{x,G}$, $\sigma_{y,G}$ | 0.25 |
| Figures 3a-d and 4a-d | $b_1$, $b_2$ | $[0.8, 1.2]$ |
| Figures 3a-d and 4a-d | $c_1$, $c_2$, $m_1$, $m_2$ | $[0.5, 1]$ |
| Figures 3a-d and 4a-d | $r_1$, $r_2$ | $[0.5, 1.5]$ |
| Figures 3a-d and 4a-d | $d$ | $[0.25, 0.55]$ |
| Figures 3a,b and 4a,b | $\tau_1$, $\tau_2$ | 0.01 |
| Figures 3c,d and 4c,d | $\tau_1$, $\tau_2$ | 1 |
| Figures 3a,c and 4a,c | $\zeta_1$, $\zeta_2$ | 0.01 |
| Figures 3b,d and 4b,d | $\zeta_1$, $\zeta_2$ | 1 |

Table B3: Figures 3 and 4 parameters. All parameters not given here are given in Table B1.
