## Appendix C for "Sick of Eating: eco-evo-immuno dynamics of predators and their trophically acquired parasites"

### Appendix C: Numerical approximation of Lyapunov exponents

Samuel R. Fleischer*[1], Daniel I. Bolnick[2], and Sebastian J. Schreiber[3]

[1] *Graduate Group in Applied Mathematics, University of California, Davis*

[2] *Department of Ecology and Evolutionary Biology, University of Connecticut*

[3] *Department of Evolution and Ecology, University of California, Davis*

May 26, 2020

Consider a system of ordinary differential equations

$$\dot{\vec{x}} = f(\vec{x}) \tag{C1}$$

and an initial condition $\vec{x}_0$ on or near an attractor of (1). Let $\vec{x}_s(t)$ be the solution of (1) with $\vec{x}_s(0) = \vec{x}_0$. Let $g(f, \vec{x}_0, t)$ denote a stable numerical algorithm that approximates the solution to (1) (i.e. Python's `scipy.integrate.odeint()`). That is,

$$g(f, \vec{x}_0, t) \approx \vec{x}_s(t). \tag{C2}$$

Let $\epsilon$ be a small positive number and choose a vector $\vec{y}_0 \neq \vec{x}_0$. For $i = 0, 1, \ldots, n$, define

$$d_i \coloneqq \|\vec{y}_i - \vec{x}_i\| \qquad \text{and} \qquad \vec{y}_i^* \coloneqq \vec{x}_i + \frac{\epsilon}{d_i}(\vec{y}_i - \vec{x}_i),$$

where $\vec{y}_i$ is defined for $i = 1, \ldots, n$ as

$$\vec{y}_i \coloneqq g(f, \Delta t, \vec{y}_{i-1}^*). \tag{C3}$$

See Figure C1 for a graphical depiction of this process.

Finally, define

$$L_i \coloneqq \ln\left(\frac{d_i}{\epsilon}\right). \tag{C4}$$

If $L_i > 0$ ($< 0$), then nearby solutions at that point move away from (towards) the reference solution $\vec{x}_s$, indicating chaotic (stable) dynamics. Then the Lyapunov exponenet $L$ for the reference trajectory $\vec{x}_s$ is defined as

$$L \coloneqq \frac{1}{n}\sum_{i=1}^{n} L_i. \tag{C5}$$

If $L < 0$, we say $\dot{\vec{x}} = f(\vec{x})$ is stable around $\vec{x}_s$ (and that $\vec{x}_s$ is a stable trajectory). If $L > 0$, we say $\dot{\vec{x}} = f(\vec{x})$ is chaotic around $\vec{x}_s$ [Sprott, 2003].

Parameters for Figure 4 of the main text are in Table B3 (Appendix B).

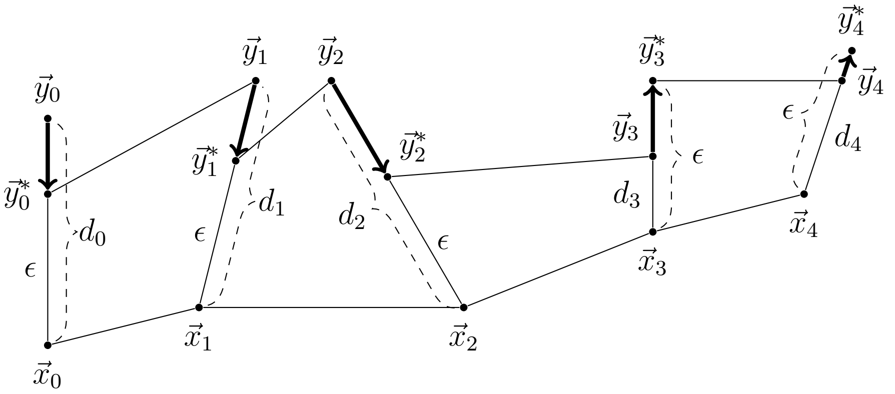


Figure C1: Cartoon of the numerical approximation of the Lyapunov exponent for the solution $\vec{x}_s$ of some system $\dot{\vec{x}} = f(\vec{x})$ with initial condition $\vec{x}(0) = \vec{x}_0$. The initial $\epsilon$-perturbation of $\vec{x}_0$ is $\vec{y}_0^*$. Each $\vec{y}_i$ is rescaled to a distance of $\epsilon$ from $\vec{x}_i$.

### References


J. C. Sprott. *Chaos and time-series analysis*, volume 69. Citeseer, 2003.
