## Appendix D for "Sick of Eating: eco-evo-immuno dynamics of predators and their trophically acquired parasites"

### Sick of Eating: eco-evo-immuno dynamics of predators and their trophically acquired parasites
### Appendix D: Determining global stability of the ecological subsystem for constant traits $\overline{x}$ and $\overline{y}$


Samuel R. Fleischer*[1], Daniel I. Bolnick[2], and Sebastian J. Schreiber[3]

[1] *Graduate Group in Applied Mathematics, University of California, Davis*

[2] *Department of Ecology and Evolutionary Biology, University of Connecticut*

[3] *Department of Evolution and Ecology, University of California, Davis*



May 26, 2020


This appendix addresses the global stability of ecological equilibria of Equation (1). In particular, for constant predator mean traits $\overline{x}$ and $\overline{y}$, we have

$$\begin{aligned} \dot{N}_1 &= N_1\left(r_1\left(1-\frac{N_1}{K_1}\right)-a_1P\right) \\ \dot{N}_2 &= N_2\left(r_2\left(1-\frac{N_2}{K_2}\right)-a_2P\right) \\ \dot{P} &= P(a_1e_1N_1+a_2e_2N_2-d) \end{aligned} \tag{D1}$$

where $a_i$ is shorthand for $\overline{a}_i(\overline{x})$ and $e_i$ is shorthand for $b_i - m_i c_i \overline{S}_i(\overline{y})$.

Note that $e_i$ may be negative. We will first discuss global stability of (D1) in the case that $e_i > 0$ for $i = 1, 2$. We will then discuss the case that $e_1 < 0$ and $e_2 > 0$ (which is symmetrical to the case that $e_1 > 0$ and $e_2 < 0$). We will conclude with the case that both $e_i < 0$ for $i = 1, 2$.

#### Case 1: Two positive conversion efficiencies

First, we nondimensionalize (D1):

$$\begin{aligned} \dot{x}_1 &= x_1(r_1 - x_1 - a_1 y) \\ \dot{x}_2 &= x_2(r_2 - x_2 - a_2 y) \\ \dot{y} &= y(a_1 e_1 x_1 + a_2 e_2 x_2 - d) \end{aligned} \tag{D2}$$

In this case, there are seven ecologically relevant equilibria of (D2):

$$\begin{aligned} (E_{+++}) &= \frac{1}{|\Theta|}(\tilde{x}_1, \tilde{x}_2, \tilde{y}), \text{ where} \\ \tilde{x}_1 &= a_1 d + a_2 e_2(a_2 r_1 - a_1 r_2) \\ \tilde{x}_2 &= a_2 d + a_1 e_1(a_1 r_2 - a_2 r_1) \\ \tilde{y} &= a_1 e_1 r_1 + a_2 e_2 r_2 - d \\ |\Theta| &= a_1^2 e_1 + a_2^2 e_2 \\ (E_{+0+}) &= \left(\frac{d}{a_1 e_1}, 0, \frac{1}{a_1}\left(r_1 - \frac{d}{a_1 e_1}\right)\right) \\ (E_{0++}) &= \left(0, \frac{d}{a_2 e_2}, \frac{1}{a_2}\left(r_2 - \frac{d}{a_2 e_2}\right)\right) \\ (E_{++0}) &= (r_1, r_2, 0) \\ (E_{+00}) &= (r_1, 0, 0) \\ (E_{0+0}) &= (0, r_2, 0) \\ (E_{000}) &= (0, 0, 0) \end{aligned}$$

The zero-species equilibrium ($E_{000}$) and the one-species equilibria ($E_{+00}$) and ($E_{0+0}$) are unstable. The two-species equilibria ($E_{+0+}$), ($E_{0++}$), and ($E_{++0}$) are globally stable if $\tilde{x}_2 < 0$, $\tilde{x}_1 < 0$, and $\tilde{y} < 0$, respectively. It is impossible for any two of $\tilde{x}_1$, $\tilde{x}_2$, and $\tilde{y}$ to be negative simultaneously. If ($E_{+++}$) is positive, then ($E_{+++}$) is globally stable. Thus in order to determine which equilibrium is globally stable, it suffices to check the signs of $\tilde{x}_1$, $\tilde{x}_2$, and $\tilde{y}$.

#### Case 2: One negative and one positive conversion efficiency

In order to keep all parameters positive, we write a new system:

$$\begin{aligned}\dot{x}_1 &= x_1(r_1 - x_1 - a_1 y)\\ \dot{x}_2 &= x_2(r_2 - x_2 - a_2 y)\\ \dot{y} &= y(-a_1 e_1 x_1 + a_2 e_2 x_2 - d)\end{aligned} \tag{D3}$$

In this case, there are six ecologically relevant equilibria of (D3):

$$\begin{aligned}(E_{+++}) &= \frac{1}{|\Theta|}(\tilde{x}_1, \tilde{x}_2, \tilde{y}), \text{ where}\\ \tilde{x}_1 &= a_1 d - a_2 e_2(a_1 r_2 - a_2 r_1)\\ \tilde{x}_2 &= a_2 d + a_1 e_1(a_2 r_1 - a_1 r_2)\\ \tilde{y} &= -a_1 e_1 r_1 + a_2 e_2 r_2 - d\\ |\Theta| &= -a_1^2 e_1 + a_2^2 e_2\\ (E_{0++}) &= \left(0, \frac{d}{a_2 e_2}, \frac{1}{a_2}\left(r_2 - \frac{d}{a_2 e_2}\right)\right)\\ (E_{++0}) &= (r_1, r_2, 0)\\ (E_{+00}) &= (r_1, 0, 0)\\ (E_{0+0}) &= (0, r_2, 0)\\ (E_{000}) &= (0, 0, 0)\end{aligned}$$

As in the previous case, the zero-species and one-species equilibria are unstable. The two-species equilibria ($E_{0++}$) and ($E_{++0}$) are globally stable if $\tilde{x}_1 < 0$ and $\tilde{y} < 0$, respectively. At this time we cannot make a statement about the global stability of ($E_{+++}$). However, if ($E_{0++}$) and ($E_{++0}$) are unstable, then the trajectory (i) approaches ($E_{+++}$) asymptotically, (ii) approaches a stable limit cycle, or (iii) is chaotic. In any case, because all solutions of (D3) are bounded, the time-average of the solution approaches ($E_{+++}$). This is the only requirement for Equation (2) to be an accurate approximation of Equation (1b) in the limit of slow evolution.

#### Case 3: Two negative conversion efficiencies

In order to keep all parameters positive, we write a new system:

$$\begin{aligned}\dot{x}_1 &= x_1(r_1 - x_1 - a_1 y)\\ \dot{x}_2 &= x_2(r_2 - x_2 - a_2 y)\\ \dot{y} &= y(-a_1 e_1 x_1 - a_2 e_2 x_2 - d)\end{aligned} \tag{D4}$$

In this case, predator fitness is always negative, and so there are only four ecologically relevant equilibria of (D4):

$$\begin{aligned}(E_{++0}) &= (r1, r2, 0)\\ (E_{+00}) &= (r1, 0, 0)\\ (E_{0+0}) &= (0, r2, 0)\\ (E_{000}) &= (0, 0, 0)\end{aligned}$$

As in the previous cases, the zero-species and one-species equilibria are unstable. The two species equilibrium ($E_{++0}$) is globally stable.
