## Appendix E for "Sick of Eating: eco-evo-immuno dynamics of predators and their trophically acquired parasites"

### Sick of Eating: eco-evo-immuno dynamics of predators and their trophically acquired parasites
### Appendix E: Unimodality condition for $\overline{W}$ with respect to $\overline{y}$

Samuel R. Fleischer*[1], Daniel I. Bolnick[2], and Sebastian J. Schreiber[3]

[1]*Graduate Group in Applied Mathematics, University of California, Davis*

[2]*Department of Ecology and Evolutionary Biology, University of Connecticut*

[3]*Department of Evolution and Ecology, University of California, Davis*

May 26, 2020

In this appendix, we assume the widths of the two susceptibility curves are equal: $\tau \coloneqq \tau_1 = \tau_2$. The predator fitness is thus

$$\overline{W} = \big(b_1 - m_1 c_1 \overline{S}_1(\overline{y})\big)\overline{a}_1(\overline{x})N_1 + \big(b_2 - m_2 c_2 \overline{S}_2(\overline{y})\big)\overline{a}_2(\overline{x})N_2 - d. \tag{E1}$$

where

$$\overline{S}_i(\overline{y}) = \beta_i - \frac{(\beta_i - \gamma_i)\tau}{\sqrt{\sigma_y^2 + \tau^2}} \exp\left[-\frac{(\overline{y} - \phi_i)^2}{2\big(\sigma_y^2 + \tau^2\big)}\right], \qquad i = 1, 2.$$

Thus, $\overline{y}$ fitness gradient is

$$\frac{\partial \overline{W}}{\partial \overline{y}} = -m_1 c_1 \overline{S}_1'(\overline{y})\overline{a}_1(\overline{x})N_1 - m_2 c_2 \overline{S}_2'(\overline{y})\overline{a}_2(\overline{x})N_2 \tag{E2}$$

$$= m_1 c_1 \left( \frac{(\beta_1 - \gamma_1)\tau(\phi_1 - \overline{y})}{(\sigma_y^2 + \tau^2)^{\frac{3}{2}}} \exp\left[ -\frac{(\overline{y} - \phi_1)^2}{2(\sigma_y^2 + \tau^2)} \right] \right) \overline{a}_1(\overline{x})N_1$$
$$+ m_2 c_2 \left( \frac{(\beta_2 - \gamma_2)\tau(\phi_2 - \overline{y})}{(\sigma_y^2 + \tau^2)^{\frac{3}{2}}} \exp\left[ -\frac{(\overline{y} - \phi_2)^2}{2(\sigma_y^2 + \tau^2)} \right] \right) \overline{a}_2(\overline{x})N_2 \tag{E3}$$

For simplicity, we introduce a composite parameter $A := \sigma_y^2 + \tau^2$ and $Z_i := \dfrac{m_i c_i (\beta_i - \gamma_i)\tau \overline{a}_i(\overline{x})N_i}{A^{\frac{3}{2}}}$ and so

$$\frac{\partial \overline{W}}{\partial \overline{y}} = Z_1(\phi_1 - \overline{y}) \exp\left[ -\frac{(\overline{y} - \phi_1)^2}{2A} \right] + Z_2(\phi_2 - \overline{y}) \exp\left[ -\frac{(\overline{y} - \phi_2)^2}{2A} \right] \tag{E4}$$

We also rescale so that $\phi_1 = 0$, $\phi_2 = \phi$, and $\overline{y} = \tilde{y}\phi$. Thus,

$$\frac{\partial \overline{W}}{\partial \overline{y}} = -Z_1 \phi \tilde{y} \exp\left[ -\frac{\tilde{y}^2 \phi^2}{2A} \right] + Z_2 \phi (1 - \tilde{y}) \exp\left[ -\frac{\phi^2 (\tilde{y} - 1)^2}{2A} \right] \tag{E5}$$

To find critical points, we set $\frac{\mathrm{d}\overline{W}}{\mathrm{d}\overline{y}} = 0$:

$$Z_1 \tilde{y} \exp\left[ -\frac{\tilde{y}^2 \phi^2}{2A} \right] = Z_2 (1 - \tilde{y}) \exp\left[ -\frac{\phi^2 (\tilde{y} - 1)^2}{2A} \right] \tag{E6}$$

Let $\Psi := \dfrac{Z_2}{Z_1} = \dfrac{m_2 c_2 (\beta_2 - \gamma_2)\overline{a}_2(\overline{x})N_2}{m_1 c_1 (\beta_1 - \gamma_1)\overline{a}_1(\overline{x})N_1}$, the ratio of the differences between maximally and minimally effective individual parasites. With this notation, we have

$$\exp\left[ -\frac{\phi^2 (\tilde{y} - \frac{1}{2})}{A} \right] = \Psi \left( \frac{1}{\tilde{y}} - 1 \right) \tag{E7}$$

This is the same form as Equation (A2) from Appendix A in Patel and Schreiber [2015]. The remainder of this appendix is a restatement of their analytical results applied to this model.

Thus, if $\Psi = 1$, then $\tilde{y} = \frac{1}{2}$ is always a critical point. If $\phi^2 < 4A$, this point is stable and if $\phi^2 > 4A$, this point is unstable. Graphical analysis shows that two additional stable equilibria exist if $\phi^2 > 4A$ (one less than $\frac{1}{2}$ and one more than $\frac{1}{2}$), and so the fitness function is bimodal when $\phi^2 > 4A$, and this corresponds to a pitchfork bifurcation.

If $\Psi \neq 1$, then the symmetry of the bifurcation breaks. If $\Psi < 1$, then parasite 1 has a greater effect on predator fitness than the parasite 2, and so we predict the critical points to be closer to $\tilde{y} = 0$ so the predator is less susceptible to limnetic parasitism. If $\Psi > 1$, we likewise expect the critical points to be closer to $\tilde{y} = 1$. We can check this by solving for $\frac{\mathrm{d}\tilde{y}}{\mathrm{d}\Psi}$:

$$\frac{\mathrm{d}\tilde{y}}{\mathrm{d}\Psi} = \frac{\dfrac{1}{\tilde{y}} - 1}{\dfrac{\Psi}{\tilde{y}^2} - \dfrac{\phi^2}{A} \exp\left[-\dfrac{\phi^2\left(\tilde{y} - \frac{1}{2}\right)}{A}\right]}. \tag{E8}$$

For all relevant values of $\tilde{y}$ ($\tilde{y} \in (0, 1)$), the numerator is positive. The denominator is positive for stable critical points and thus $\frac{\mathrm{d}\tilde{y}}{\mathrm{d}\Psi} > 0$ for stable critical points in $(0, 1)$. Thus, the stable phenotype values of $\tilde{y}$ increase as $\Psi$ increases.

### References

S. Patel and S. J. Schreiber. Evolutionarily driven shifts in communities with intraguild predation. *The American Naturalist*, 186(5):E98–E110, nov 2015. doi: 10.1086/683170. URL `https://doi.org/10.1086/683170`.
